# Investigating the effect of channel pruning on functional near-infrared spectroscopy data collected from children aged 5-24 months

**DOI:** 10.1101/2025.10.06.680288

**Authors:** Samuel Beaton, Borja Blanco, Chiara Bulgarelli, Clare Elwell, Sarah Lloyd-Fox, Ebrima Mbye, Samantha McCann, Anna Blasi Ribera, Sophie Moore

## Abstract

**Significance:** Infant functional near-infrared spectroscopy (fNIRS) data are particularly vulnerable to noise; participant behaviour can result in motion artefacts and reduced set-up times can cause poor optode coupling. Accurate channel pruning is therefore essential but approaches vary and often use adult-derived thresholds, risking unnecessary data loss.

**Aim:** This work systematically compared pruning approaches and parameter choices to evaluate their effects on data quality and retention in infant fNIRS.

**Approach:** Data from 5–24 month-old infants were collected across two cohorts, using two paradigms. Channel pruning was performed using the coefficient of variation (CV) and the Quality Testing of Near Infrared Scans (QT-NIRS) tool, varying key thresholds. Multilevel models assessed effects of pruning method, parameter choice, age, motion, and testing site on signal-to-noise ratio (SNR) and channels retained.

**Results:** QT-NIRS produced significantly higher SNR than CV pruning across nearly all age, task, and cohort combinations, when matched for data retention. Higher QT-NIRS thresholds improved quality but reduced retention. Motion prevalence strongly reduced both SNR and retention; testing site and age had smaller but notable effects.

**Conclusions:** QT-NIRS offers a better balance of data quality and retention than CV pruning. Lower QT-NIRS thresholds than adult defaults are recommended for infant data. These findings provide practical guidance for preprocessing pipelines in developmental fNIRS research.

## 1 Introduction

### 1.1 Scalp-optode coupling

In a typical functional near-infrared spectroscopy (fNIRS) experiment, participants wear a headband or cap embedded with optodes to monitor brain activity by emitting and detecting near-infrared light at two separate wavelengths. When fitted securely, the cap ensures a motion-robust signal, enabling the study of a wide range of cognitive tasks and abilities with less stringent demands for stillness than other neuroimaging modalities [1]. For this reason, fNIRS is widely used with developmental populations [2]. However, signal processing is usually still required to remove artefacts arising from motion, poor scalp-optode coupling, and physiological signal confounds [3], [4], [5]. Infant data is particularly susceptible to motion and poor scalp-optode coupling, as infants typically exhibit increased fussiness, limited compliance with instructions, and shorter attention spans, which increase the likelihood of participant motion, reduced capping time and difficulties in handling the imaging headgear [6]. In fact, poor coupling allows light from the source optode(s) to escape, or ambient light to flood detector optode(s) [7]. Affected channels can exhibit signal saturation (easily detectable by unrealistically high raw intensity values caused by excessive light reaching the detector) and greater variability, which impacts the estimation of the haemodynamic response [8] whose amplitude is already lower and more variable in infants than in older participants [9], [10].

### 1.2 Strategies to mitigate poor optode coupling

Fitting the fNIRS headgear securely aids scalp-optode coupling [7] but is time-consuming and assumes stable coupling throughout recording, which is challenging when working with infant participants. To reduce the impact of poor coupling on data quality, post-hoc channel pruning – the exclusion of data from an entire channel – is therefore often required. An important pruning consideration is the trade-off between data retention and quality: removing poorly-coupled channels improves overall signal quality and mitigates the impact of superfluous signals, but reduces the number of remaining channels and participants available for analysis [6]. This is particularly important in infant research given the already high attrition rates due to low attention span and susceptibility to fussiness [11]. Pruning method selection is important, and manually pruning channels is subjective and time-consuming [8], especially for high-density imaging arrays. Two methods are often used to prune channels: the coefficient of variation (CV) or the ‘Quality Testing of Near-InfraRed Spectroscopy’ (QT-NIRS) tool.

CV pruning quantifies the relative signal variability. Channels are pruned if the CV for either wavelength falls below a particular threshold [12], [13], if the CV difference between wavelengths exceeds a threshold [14], or both [15]. While this method is faster and less subjective than manual pruning, a parameter choice is still required and some levels of variability - which is expected in the task-based fNIRS signal due to the evoked haemodynamic response [16] - may be interpreted as signal noise. Additionally, CV pruning only examines the signal in the time domain, potentially overlooking important signal properties.

QT-NIRS utilizes objective signal measures from both the time- and frequency domains [7], [16], providing a more comprehensive consideration of signal quality. Channels are pruned via signal characteristic assessments in two domains using the scalp coupling index (SCI) and peak spectral power (PSP). The SCI is a time-domain approach, which assesses correlation between wavelengths within the cardiac frequency band after bandpass filtering the signal, with high values indicative of a strong cardiac component [17]. This incorporates a time-domain signal characteristic, but also risks retaining channels with high correlation due to motion-induced artefacts. To address this, QT-NIRS incorporates a second measure based on the frequency domain, PSP, which detects strong, recurrent oscillations in the signal. High PSP values in the cardiac frequency band likely correspond to cardiac pulsations, whereas components with inconsistent or varying frequencies usually result in lower PSP values. QT-NIRS utilizes strengths from manual pruning (physiological grounding, consideration of both time- and frequency domains) and CV pruning (objectivity; efficiency) making it a robust tool for assessing data quality at the channel level.

Recently, QT-NIRS has been increasingly adopted in infant fNIRS research [18], [19] yet independent comparisons with other pruning approaches are yet to be established, and empirical estimations of SCI and PSP parameters are available only for adult participants [7], [16]. This highlights the need to refine its implementation with infant data. In addition to behaviour, both physiological and anatomical factors may affect channel pruning in infants: they have thinner scalp tissue and higher cardiac signal frequencies (∼1.3–3.2 Hz at rest, compared to ∼1–1.7 Hz in adults)[20], [21]. The former may result in a weak superficial cardiac signal, whereas the latter may result in coarse representations of the infant cardiac signal by fNIRS instrumentation sampling rates optimized for adult participants [22]. Further, signal quality can be detrimentally affected by skin and hair colour, hair type, age and even head size with darker skin pigmentation and thicker hair corresponding to poorer signal quality when compared to other skin and hair types [23], illustrating the need for inclusion of less-frequently sampled populations in fNIRS studies.

Against this backdrop, the objectives of this work are to:

(a) Compare QT-NIRS as a pruning method against pruning using CV, which is used frequently by fNIRS users – and provides a baseline for channel pruning via a previously employed method and parameters
(b) Investigate contextual and data-derived measures which affect data quality and channel retention (with a particular focus on QT-NIRS parameter choices, age and motion incidence)
(c) Provide guidance on channel pruning and QT-NIRS use for infant participants

To achieve this, fNIRS data from the Brain Imaging for Global HealTh (BRIGHT) Project [24], a longitudinal study of infant development in Kiang West (The Gambia) and Cambridge (UK), were analyzed. The analyses in this work incorporates data from two different experimental paradigms, collected from both the Gambian and UK sites and pertaining to participants with physical and behavioural characteristics from both a commonly-sampled and a more underrepresented population in fNIRS research [23]. The longitudinal nature of the data (with five time points over the first two years of life) further enables the investigation of the effects of age across early childhood while accounting for variability in cohort and task.

Based on prior literature and exploratory findings, the following hypotheses were formulated:

(1) QT-NIRS based pruning will result in a greater balance of data quality and retention compared to CV, due to its multidomain examination of signal characteristics
(2) Motion will be negatively associated with signal quality and retention, because of (i) increased likelihood of latent, undetected artefacts in data where more motion is detected, (ii) displacement of optodes affecting signal quality after motion artefacts, or both.
(3) Data quality and retention will be diminished in the Gambian cohort, given the increased probability of darker, coarser hair types interfering with optode-scalp coupling

## 2 Methods

### 2.1 Data

#### 2.1.1 Participants

Participants were recruited into the BRIGHT project, from early 2016 to February 2018, and fNIRS data were collected when infants were 1-, 5-, 8-, 12-, 18- and 24 months (hereafter *x*mo for *x* months of age), plus a follow-up between 3 and 5 years of age [25], [26]. The 1mo fNIRS protocol was limited to auditory stimuli with sleeping participants [27], likely inducing infant motion with a different noise profile; at 3-5 years, a different fNIRS cap for data collection was used and data at this age were not collected in the UK site. To enable matched dataset comparisons, data from the other 5 time points (5-, 8-, 12-, 18- and 24mo) was therefore used in this work. Participants met the inclusion criteria if infants were born at 37–42 weeks’ gestation (both cohorts) and had a minimum birth weight of 2.5 kg (UK only).

After applying exclusion criteria, a total of 204 mother-infant dyads were included in the Gambian cohort; of these, 185 remained at the 24mo timepoint. Pregnant, Mandinka-speaking women were recruited during routine antenatal clinical assessments at MRCG@LSHTM Keneba Field Station by fieldworkers in the Gambian BRIGHT Project team. An information sheet and consent form written in English were provided to potential recruits then explained fully in Mandinka by a study staff member. fNIRS data collection took place at MRC Unit The Gambia at the London School of Hygiene and Tropical Medicine (‘MRCG@LSHTM’) Keneba Field Station. Ethical approval was granted by the joint Gambia Government/MRC Ethics Committee under the title: ‘Developing brain function for-age curves from birth using novel biomarkers of neurocognitive function’, SCC number 1451v2.

61 mother-infant dyads were enrolled in the UK from the Rosie Hospital, Cambridge University Hospitals NHS Foundation Trust. Information about BRIGHT was provided during antenatal appointments, with families expressing an interest contacted and recruited subsequently via email or phone call. Data collection primarily took place at Evelyn Perinatal Imaging Centre at Rosie Hospital, Addenbrooke’s Hospital, Cambridge, and to a lesser extent at the Centre for Brain and Cognitive Development in Cambridge [24], [28]. Ethical approval was given by the National Research Ethics Service Committee East of England (REC reference 13/EE/02000); informed written consent was obtained from all parents/carers prior to participation.

#### 2.1.2 fNIRS Paradigms

##### Social/Non-Social paradigm

Full details of the Social/Non-Social (SNS) task paradigm can be found in [15] and [29]. Briefly, the paradigm consisted of alternating visual social (silent), auditory social and auditory non-social stimuli. Stimuli repeated until the participant became bored or fussy or the end of the task was reached; inter-stimulus baselines varied between 9 to 12 seconds.

Visual social stimuli consisted of full colour, life-sized videos of adults from the same population as the participant, on a 24-inch screen ∼100cm away. Throughout, adult actors in the video either moved their eyes or played ‘hand-games’ for 9-12 seconds. Actors, their actions, and concurrent spoken auditory stimuli were varied to prevent anticipatory brain activity. Auditory stimuli were non-synchronized to the video in terms of both duration (8 seconds) and content, with environmental sounds for non-social stimuli and non-vocal speech sounds for social stimuli. Sounds in each condition (social and non-social) were matched for duration and sound intensity.

##### Habituation and novelty detection (HaND) paradigm

The experimental paradigm included 25 trials, each consisting of a spoken 8sec sentence in the family’s first language (i.e. English or Mandinka) followed by 10sec of silence. The first trial was preceded by at least 10sec of silence, acting as a baseline.

The same recording, with a female voice, was used for trials 1-15; a different, male voice was used for trials 16-20; finally, the original, female recording was again used for trials 21-25. The stimulus sentence: *“Hi baby! How are you? Are you having fun? Thank you for coming to see us today. We’re very happy to see you”* was translated to Mandinka to maintain the same semantic meaning.

Technical detail on the recording, processing and playback of the auditory stimulus can be found in previous work [14], [26].

#### 2.1.3 Data collection

Custom-made headgear was fitted after head measurements (head circumference, and ear-to-ear both around the forehead and over the top of the head) had been taken to aid with the alignment of fNIRS headgear with the 10/20 system anatomical landmarks. Headgear consisted of custom-built stretchy silicone headbands to increase friction and prevent slippage, with attached probes into which optodes were clipped. Optodes were designed to accommodate glass optic fibres at right-angles to allow them to sit flush on the scalp. The headband was fastened around the head to provide even pressure over the base of the probes [30], [31]. In the left hemisphere, headgear was placed such that source 4 in Figure 1 was centered above the preauricular point, so that the channel it formed with the detector located directly behind it sat above T3 in the 10-20 system; the equivalent right hemisphere channel was above T4. The array angle was guided by the headband, which was placed on the head so that it touched the join between the ear and head and, frontally, lay over the infant brow line (through Fp1 and Fp2 in the 10-20 system) [32].

**Figure 1:**
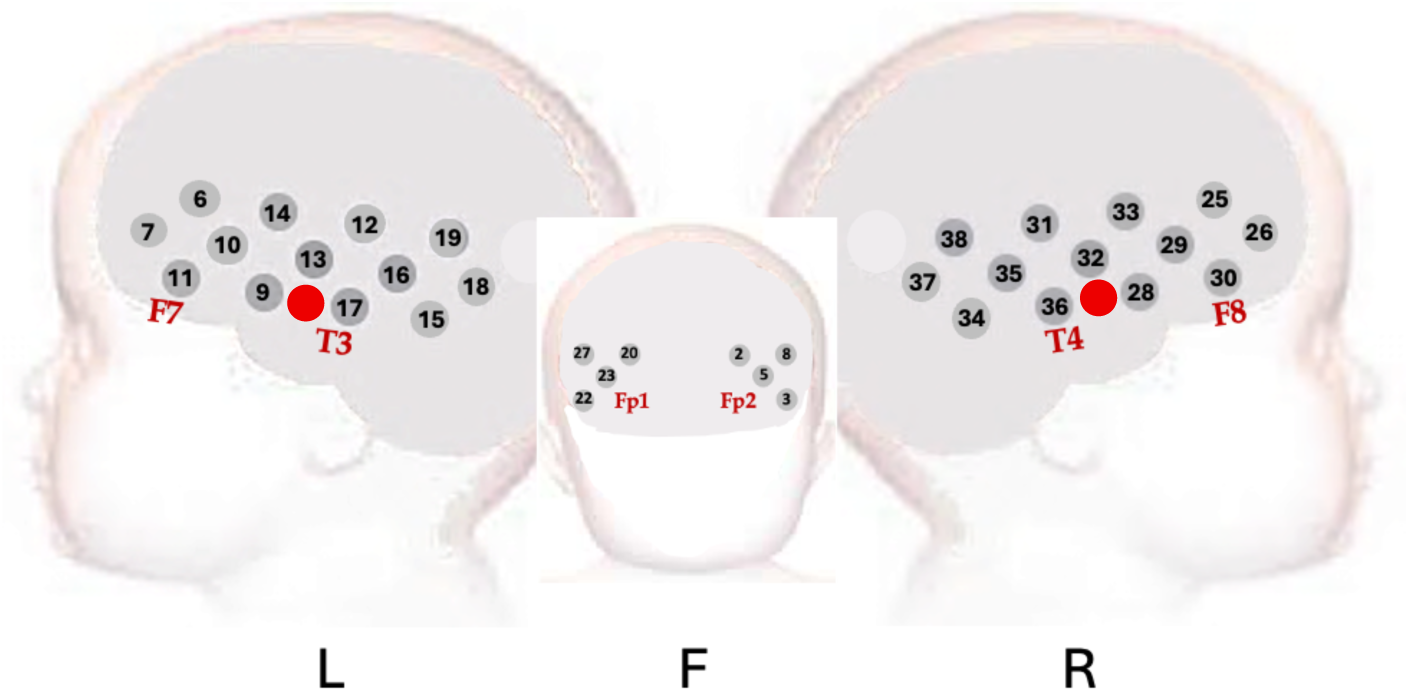
Array layout during data collection. Red dots indicate position of optodes centred above the preauricular point during headgear fitting.

The headgear was designed to record responses bilaterally from auditory-associative brain regions, including the inferior frontal gyrus (IFG), middle and superior temporal regions, and the temporo-parietal junction [14], [33], [34]. The fNIRS array comprised 17 channels in each hemisphere with a source-detector (S-D) distance of 2cm, corresponding to a penetration depth of ∼1cm from the skin surface and thus permitting measurement of the gyri and superficial sulci [14]. fNIRS data were collected using this array design with the NTS optical topography system (Gowerlabs Ltd., UK) with a sampling frequency of 10Hz and source wavelengths of 850 and 780nm.π

Infants sat on a carer’s lap during data acquisition. Carers were discouraged from interacting with the infant to attempt to minimize confounding stimuli however infants’ attention was engaged, if necessary, with (non-social, non-auditory) bubble-blowing and silent demonstration of soft toys, which also minimized infant head movement. The HaND task was part of a larger battery of fNIRS assessments with a total recording time of ∼ 21min 30s (6min social task; 4min functional connectivity data acquisition; 7min 30s HaND; 4min further functional connectivity). Where possible, paradigms were completed uninterrupted; sessions were paused and subsequently resumed in the event of infant discomfort.

### 2.2 Channel Pruning

#### 2.2.1 Pre-pruning steps

First, channels were excluded from analyses if their minimum light intensity value was less than 3e-4, based on previous experience with the NTS system [14]; these were labelled ‘channels with signal extrema’ (CSE). Analysis of the pruning methods was conducted on motion-free segments. To find motion-free data, motion artefacts were detected using *hmrMotionArtifactByChannel* function from Homer2 [36] with established infant fNIRS preprocessing parameters: *tMotion = 1, tMask = 1, STDEVthresh = 15* and *AMPthresh = 0.4* [37]. Data for each channel was split into 3 second windows per channel, as this aligned with QT-NIRS temporal segmentation and thus avoided additional processing complexities. Windows were excluded from pruning analyses if they contained artefacts at any point. If the data for a particular channel at one wavelength was excluded, data for both wavelengths was removed.

The two channel pruning methods assessed in this work were implemented using custom-written scripts in MATLAB [26]. Both pruning methods described use raw light intensity data as input.

#### 2.2.2 CV Pruning

CV pruning was conducted using an in-house script, and CV itself was calculated on motion-free data for each wavelength and channel using the equation:

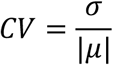

where *σ* and *μ* are the standard deviation and mean of the light intensity signal in a channel, respectively [38]. Channels were pruned if the difference between CV values for each wavelength exceeded 0.2 based on previous experience with the same data and system [14].

#### 2.2.3 QT-NIRS pruning

QT-NIRS was implemented using the function *qtnirs* (available at https://github.com/lpollonini/qt-nirs at the time of writing) for precise control over the pruning and quick, repeated processing of the large volume of data.

More detail on QT-NIRS can be found in publications describing the methods [7], [17] but an outline is provided here. First, bandpass filtering is conducted to retain only those frequencies in the cardiac band, ∼1.3 − 3.2*Hz* in the case of infants [20]. The cross-correlation of contemporaneous (zero-lag) wavelength signals in the cardiac frequency band, (i.e., SCI), is then calculated:

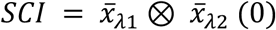

where *x̄*_*λi*_ represents the light intensity signal for wavelengths *i* = 1, 2 in motion-free signal. PSP is the maximum signal value in the frequency domain, representing the dominant oscillation in the bandpass-filtered signal and presumed to correspond to the cardiac frequency in well-coupled data. The recorded signal is divided into equal-length windows, and both SCI and PSP are calculated for each window. A window’s signal is considered of sufficient quality if both the calculated SCI and PSP exceed the user-defined thresholds, *sci_threshold* and *psp_threshold*, respectively.

The focus was on the alteration of *sci_threshold* and *psp_threshold* during analysis, since each provides a threshold for one of the key measures of optode coupling quality used to assess data quality with QT-NIRS, and adult reference values of *sci_threshold* = 0.8 and *psp_threshold* = 0.1 are available for these two parameters [16]. Default parameters were used for window size (3 seconds) and quality threshold, or *q_threshold* (0.75), which prunes channels with less than 75% of windows meeting both SCI and PSP threshold values.

### 2.3 Statistical analyses

Multi-level models (MLMs), a form of linear regression that estimates variance at multiple levels, were used for statistical analyses, to effectively account for repeated measures, hierarchical data structures (including participant- and channel-level measures), and missing data [39]. All models were fitted in R 4.4.1 [40] using the *lme4* package [41]. All models included random intercepts for each participant to account for individual variation. Final models used for analysis are described in ‘Models and Analyses’.

#### 2.3.1 Measures, Outcomes and Effects

Considering the anticipated data quality/inclusion trade-off, which is central to decision making around preprocessing of fNIRS data [6], the performance of pruning methods and parameters were assessed using two metrics: (i) signal-to-noise ratio (SNR) and (ii) channel inclusion/exclusion percentage.

The effects of other factors such as age, motion, and signal extrema, were included as predictors. Other measures were more particular to this dataset - such as task, cohort and optode position. They were included as covariates to account for the variability they may cause.

The variables used are listed below, with letters contained in brackets indicating whether they were used as outcome variables (O), predictors (P), or covariates (C). Full rationale can be found in Supplementary Materials 1.

##### 2.3.1.1 Task-relevant channel signal-to-noise ratio (O)

Signal quality after pruning was measured on the included channels using the signal-to-noise ratio (SNR):

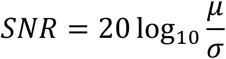

with *μ* and *σ* the mean and standard deviation of the signal, respectively [5]. SNR was calculated in ‘task-relevant channels’ (TRCs) – channels where a true haemodynamic signal was observed. Age-specific TRCs were determined using prior analyses: the SNS TRCs were taken from work by Benerradi and colleagues [25] whereas the HaND TRCs were taken from work by Blasi and colleagues [26]. TRCs for each age and task were those which exhibited a haemodynamic response in both chromophores for either cohort, except the SNS task at 18mo: only 1 channel met this criterion, so channels were added for this age/task combination if they were in the set of TRCs for at least two other ages for the SNS task. This outcome was named the task-relevant channel signal-to-noise ratio (TRC SNR).

##### 2.3.1.2 Channels retained (O)

The number of channels per participant included after pruning using the described method and parameter(s) was summed – this was labelled ‘Channels Retained’ (CR).

##### 2.3.1.3 SCI Threshold (P)

*sci_threshold* values ranging from 0.05 to 0.9 were used, with increments of 0.05. The upper threshold of 0.9 sits between the recommended value for adult participants (0.8) and the theoretical maximum of SCI = 1. Initial exploratory analyses (see Supplementary Materials 2) for each age/task/cohort combination indicated that even very low SCI values continued to alter signal quality, so the entire range of lower values was used.

##### 2.3.1.4 PSP Threshold (P)

The *psp_threshold* value was varied, ranging from 0.005 to 0.1, with increments of 0.005. The upper value of 0.1 is the recommended threshold for adult participants. Values which spanned the entire possible range lower than this – based on initial results from simpler MLM analyses – were used; this is supported by the considerable proportion of infant PSP values lower than the adult threshold apparent during post-analysis data examination in the UK cohort (see Figure 2).

**Figure 2:**
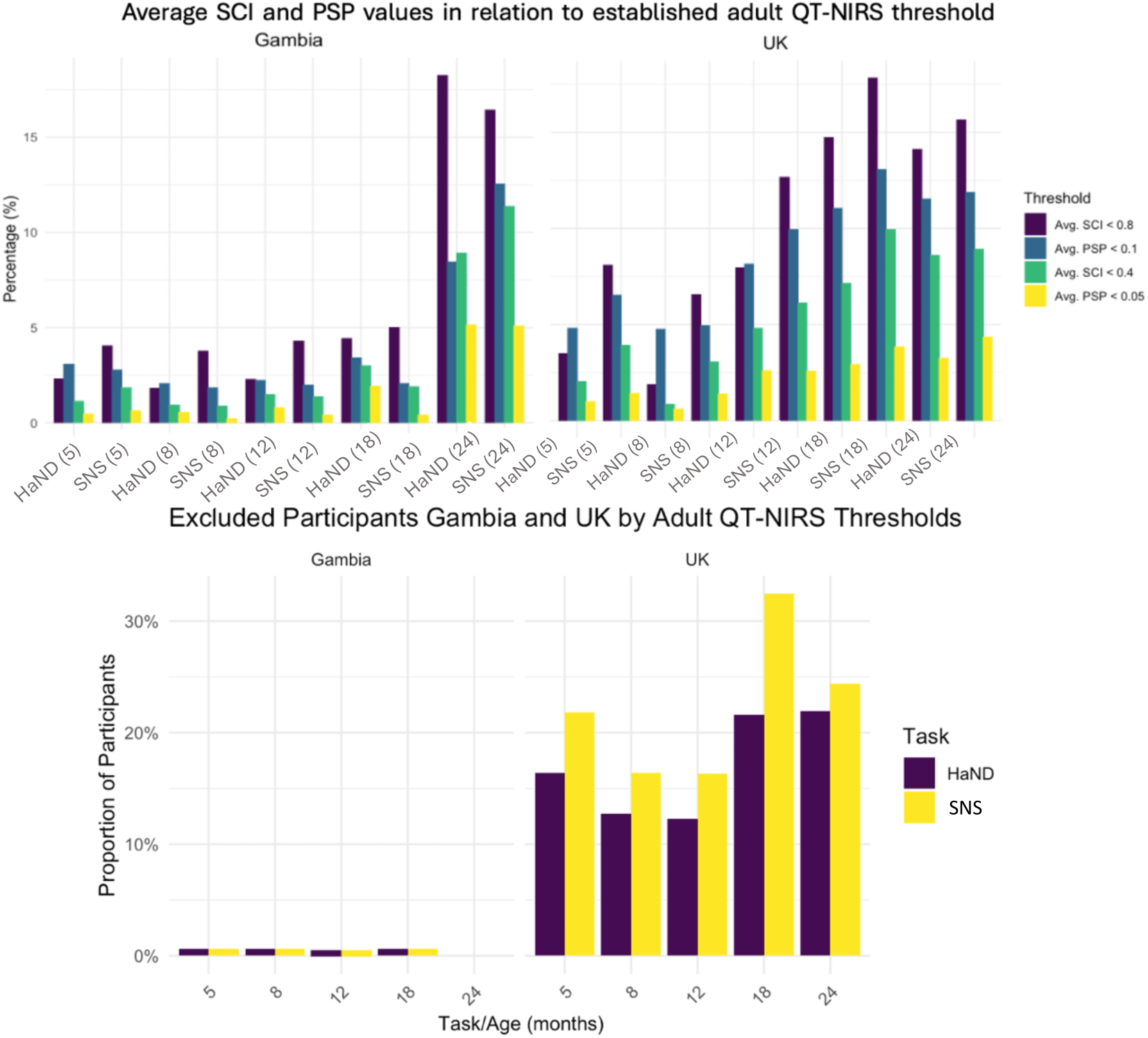
Data characteristics in relation to adult QT-NIRS thresholds. Top: bar charts showing the proportion low channel Average SCI and Average PSP measures in relation to the adult recommended parameters of 0.8 (SCI) and 0.1 (PSP), and half of these threshold values (0.4 and 0.05, respectively). Top-left: Gambia cohort. Top-right: UK cohort. Bottom: number of participants, by age and Task, with less than 60% of acceptable channels exhibiting mean SCI and PSP values compared to adult threshold values of 0.8 and 0.1, respectively. Highest exclusion rate was ∼33% for UK infants at 18mo during the SNS task.

##### 2.3.1.5 Percentage of motion (P)

For each participant/age/task combination, the cross-channel mean of the percentage of windows per channel containing motion as identified by *hmrMotionArtifactByChannel* and subsequently excluded from analyses (see ‘Pruning’) was labelled the ‘Percentage of Motion’ (PoM) providing a measure of the prevalence of motion.

##### 2.3.1.6 Age (P)

Age was included as a five-level predictor (5-, 8-, 12-, 18-, 24mo) to assess change with age.

##### 2.3.1.7 Cohort (C)

Cohort was included as a two-level covariate (Gambia and UK).

##### 2.3.1.8 Task (C)

Task was included as a two-level covariate (HaND and SNS) to account for the contribution to variance of the different paradigms.

##### 2.3.1.9 Channels pruned due to signal extrema (C)

The number of CSE per participant, for each age and task, was used as a participant-level covariate and proxy measure of poor optode coupling.

#### 2.3.2 Models and analyses

##### 2.3.2.1 Comparison of QT-NIRS and CV pruning

In line with objective (a), each fNIRS recording underwent channel pruning using:

1. CV, by pruning channels where the CV values for different wavelength signals differed by 20% or more (‘CV’)
2. *sci_threshold* only at every parameter value, by setting *psp_threshold* to 0, (‘SCI Only’), and
3. every combination of *sci_threshold* and *psp_threshold* parameters listed in ‘Measures, Outcomes and Effects’ (‘Full QT-NIRS’)

Pruning method 1 provides a baseline for the channel pruning, using a method and pruning criteria which has previously been used for the HaND task [14]. Pruning approach 2 permits comparison of two time-domain methods (methods 1 and 2) to assess the impact of incorporating temporally specific measures. Pruning approach 3 provides insight into the benefit of additionally using a PSP threshold and pruning using frequency characteristics of the signal.

For every age/task/cohort combination, each *sci_threshold* parameter choice (SCI) or combination of *sci-* and *psp_threshold* values (QT-NIRS) were given two separate rankings according to their similarity to CV pruning, in terms of the mean number of channels retained across all participants and the total number of participants excluded. These two rankings were then combined to find the parameter (SCI) or parameters (QT-NIRS) producing the closest approximation to data retention provided by CV pruning.

For each of the 20 age (5 levels)/task (2 levels)/cohort (2 levels) combinations, the following model was then fitted to participant-level data:

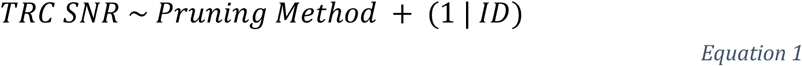

to compare the effect of pruning methods (1)-(3) on signal quality whilst using a random intercept for each infant to account for inter-participant variability. To correct for multiple comparisons, *p*-values were Bonferroni-corrected.

##### 2.3.2.2 Effect of SCI and PSP threshold choice on signal quality and retention

Models were designed to investigate the effect of *sci_threshold*, *psp_threshold*, age, and motion on TRC SNR and Channels Retained (Objectives (b) and (c)).

To examine potential combinations of theoretically viable predictors, covariates, and their interactions used for each outcome, a systematic approach to model building was used as a first step, combining predictors in various model formulae using combinatorial logic before assessing model fit. Model fit was assessed using the Akaike Information Criterion (AIC) [42], a model selection criterion that balances goodness of fit with model complexity [43, p. 824]. Model variables were also included based on the outcomes of the subsidiary investigations described in Supplementary Materials 3 which investigated the factors affecting motion incidence and average SCI and PSP measures at the channel level. These additional variables are included in the bottom line of Equation 2.

Mindful of model convergence issues and overfitting, the priority when constructing models was to include predictors of interest, plus interaction terms, random slopes and random intercepts which incorporated them. This resulted in the following model for both outcomes TRC SNR and Channels Retained:

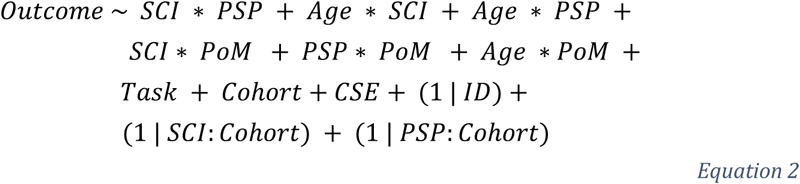

where SCI = *sci_threshold*, PSP = *psp_threshold*, and symbol ‘∗’ denotes an interaction term (itself denoted using ‘:’) plus all individual terms used in the interaction, as is consistent with notation used in the *lme4* package. Random intercept terms (bottom row) were included due to quantitative support based on subsidiary MLM investigations into average SCI and PSP measures.

Assessments of the model residuals showed that they were non-normally distributed, so to calculate effects a bootstrapping approach was used. Bootstrap datasets were generated by sampling rows from the original dataset with replacement, using a fixed random seed for reproducibility. The relevant model for each bootstrap sample was fitted using the *lmer* function from the *lme4* package and extracted fixed effect estimates. To mitigate potential biases caused by fitting models to data with different scales, variables *x*_*i*_ were scaled and centred:

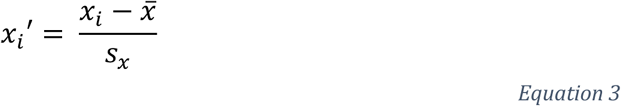

where *x*_*i*_′ is the scaled value, and *x̄* and *s*_*x*_ are the sample mean and standard deviation, respectively, of variable *x*.

## 3 Results

CV pruning yielded an average TRC SNR of 23.7 ± 1.66 across all 20 age/task/cohort combinations. Pruning with QT-NIRS using just the SCI threshold resulted in a mean increase of 2.13 ± 0.644 in TRC SNR beyond that obtained using CV pruning. Similarly, using both SCI- and PSP thresholding in combination resulted in a mean TRC SNR increase of 2.16 ± 0.627 in comparison to CV pruning (see Figure 3, and Supplementary Materials 4 for the full comparison of TRC SNR values).

**Figure 3:**
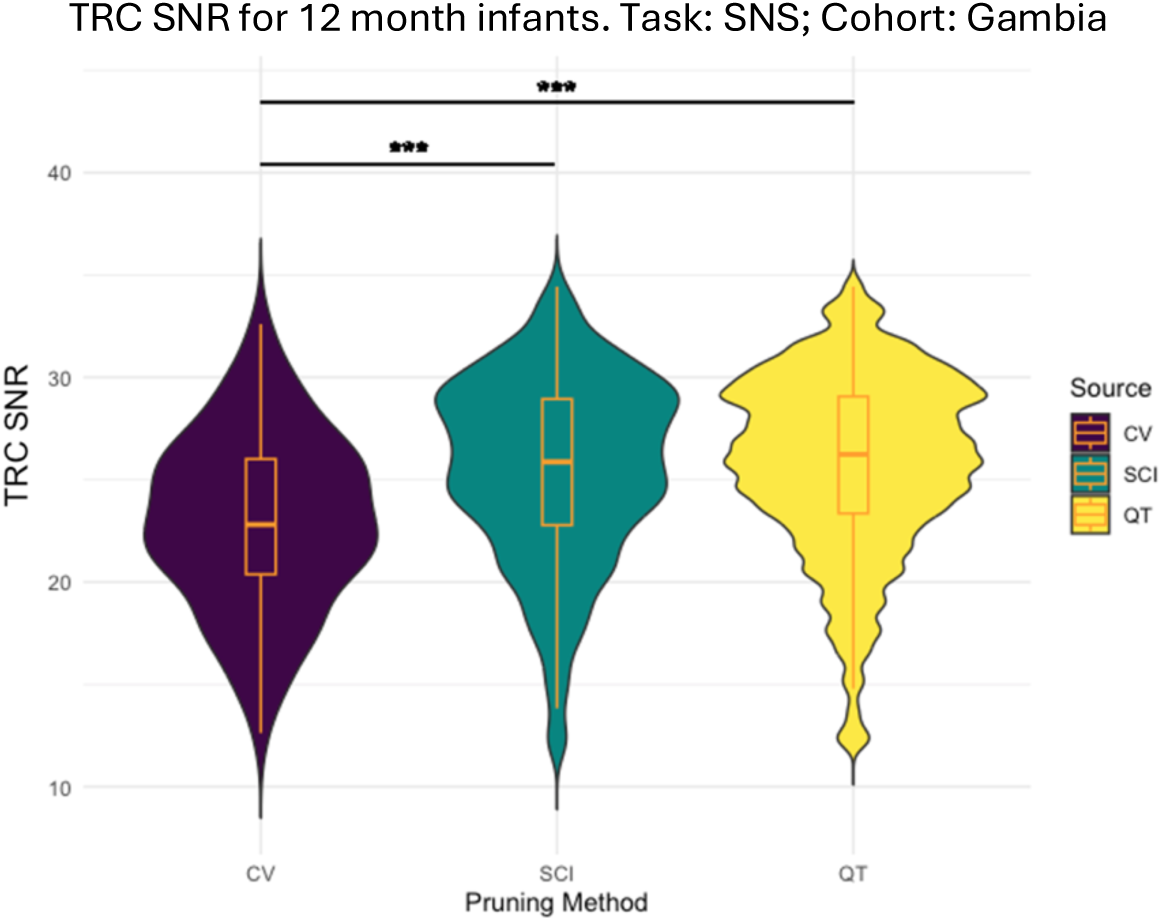
Example violin plot demonstrating typical differences between obtained mean TRC SNR values using CV pruning, QT-NIRS using SCI threshold only, and QT-NIRS utilising both parameters for Gambian participants at 12mo during the SNS task.

### 3.1 Comparison of QT-NIRS and CV pruning

#### 3.1.1 Pruning using CV and QT-NIRS

The effect size and Bonferroni-corrected significance values of the 3 contrast conditions were calculated using Equation 1; results are displayed in **Table 1**. For 19 of the 20 age/task/cohort combinations, significant (p < 0.01) positive effects were found on the TRC SNR when controlling for data rejection when using QT-NIRS, using either the SCI Only or Full QT-NIRS approach. In 17 of 20 combinations, the differences were found to be statistically significant at the p < 0.0001 threshold. Though the effect was positive for the single remaining combination of 20, it did not reach statistical significance.

**Table 1:** Condition contrasts between each of the three pruning methods.

| Age (months) | Task | Cohort | CV vs SCI Only |  | CV vs Both Parameters |  | SCI Only vs Both Parameters |  |
| --- | --- | --- | --- | --- | --- | --- | --- | --- |
|  |  |  | Significance | Effect Size | Significance | Effect Size | Significance | Effect Size |
| 5 | HaND | Gambia | <0.001 | 20.85 | <0.001 | 20.83 | 1.0 | 0.05 |
| 8 | HaND | Gambia | <0.001 | 12.71 | <0.001 | 12.75 | 1.0 | <0.01 |
| 12 | HaND | Gambia | <0.001 | 19.32 | <0.001 | 19.70 | 1.0 | 0.27 |
| 18 | HaND | Gambia | <0.001 | 17.04 | <0.001 | 17.10 | 1.0 | 0.04 |
| 24 | HaND | Gambia | <0.001 | 12.36 | <0.001 | 12.12 | 1.0 | 0.17 |
| 5 | SNS | Gambia | <0.001 | 16.97 | <0.001 | 16.93 | 1.0 | 0.02 |
| 8 | SNS | Gambia | <0.001 | 14.69 | <0.001 | 14.70 | 1.0 | 0 |
| 12 | SNS | Gambia | <0.001 | 16.95 | <0.001 | 17.65 | 1.0 | 0.49 |
| 18 | SNS | Gambia | <0.001 | 15.10 | <0.001 | 15.09 | 1.0 | <0.01 |
| 24 | SNS | Gambia | <0.001 | 13.38 | <0.001 | 13.38 | 1.0 | 0 |
| 5 | HaND | UK | <0.001 | 9.40 | <0.001 | 9.40 | 1.0 | <0.01 |
| 8 | HaND | UK | <0.001 | 12.90 | <0.001 | 12.90 | 1.0 | <0.01 |
| 12 | HaND | UK | <0.001 | 8.70 | <0.001 | 8.56 | 1.0 | 0.10 |
| 18 | HaND | UK | <0.001 | 10.92 | <0.001 | 10.92 | 1.0 | <0.01 |
| 24 | HaND | UK | 0.012 | 4.68 | 0.012 | 4.68 | 1.0 | 0 |
| 5 | SNS | UK | <0.001 | 6.93 | <0.001 | 7.80 | 1.0 | 0.62 |
| 8 | SNS | UK | <0.001 | 7.18 | <0.001 | 7.28 | 1.0 | 0.07 |
| 12 | SNS | UK | 0.924 | 3.23 | 0.288 | 3.87 | 1.0 | 0.45 |
| 18 | SNS | UK | 1.0 | 2.84 | 1.0 | 2.84 | 1.0 | 0 |
| 24 | SNS | UK | 1.0 | 0.76 | 1.0 | 0.76 | 1.0 | <0.01 |

Methods are: CV, QT-NIRS using the *sci\_threshold* parameter only, and QT-NIRS using both the *sci\_threshold* and *psp\_threshold* parameters. Significance values calculated during bootstrapping and reported after correction. Effect sizes rounded to 2d.p.

#### 3.1.2 Pruning using SCI Only compared with both parameters

No significant statistical differences were found between TRC SNR values when pruning using SCI only and Full QT-NIRS approaches. In every case, however, the mean TRC SNR across participants was higher when using both parameters than when using *sci_threshold* alone.

A representative comparison of the three different pruning methods is given in Figure 3, with higher average TRC SNR values obtained using both QT-NIRS approaches when compared to CV pruning. A higher mean TRC SNR is obtained using both parameters when pruning with Full QT-NIRS compared to using SCI Only, but with a more erratic distribution of values.

### 3.2 Predictors of TRC SNR and Channels Retained

The focus here is primarily on reporting results for the outcomes of interest and fixed effects with wider generalisability but full results are included in Figure 4. Effect sizes were classified as small, medium or large if the absolute value of the Estimate was < 0.15, < 0.35, or ≥ 0.35, respectively, using Cohen’s *f*^2^thresholds (Cohen, 2013, chap.9).

**Figure 4:**
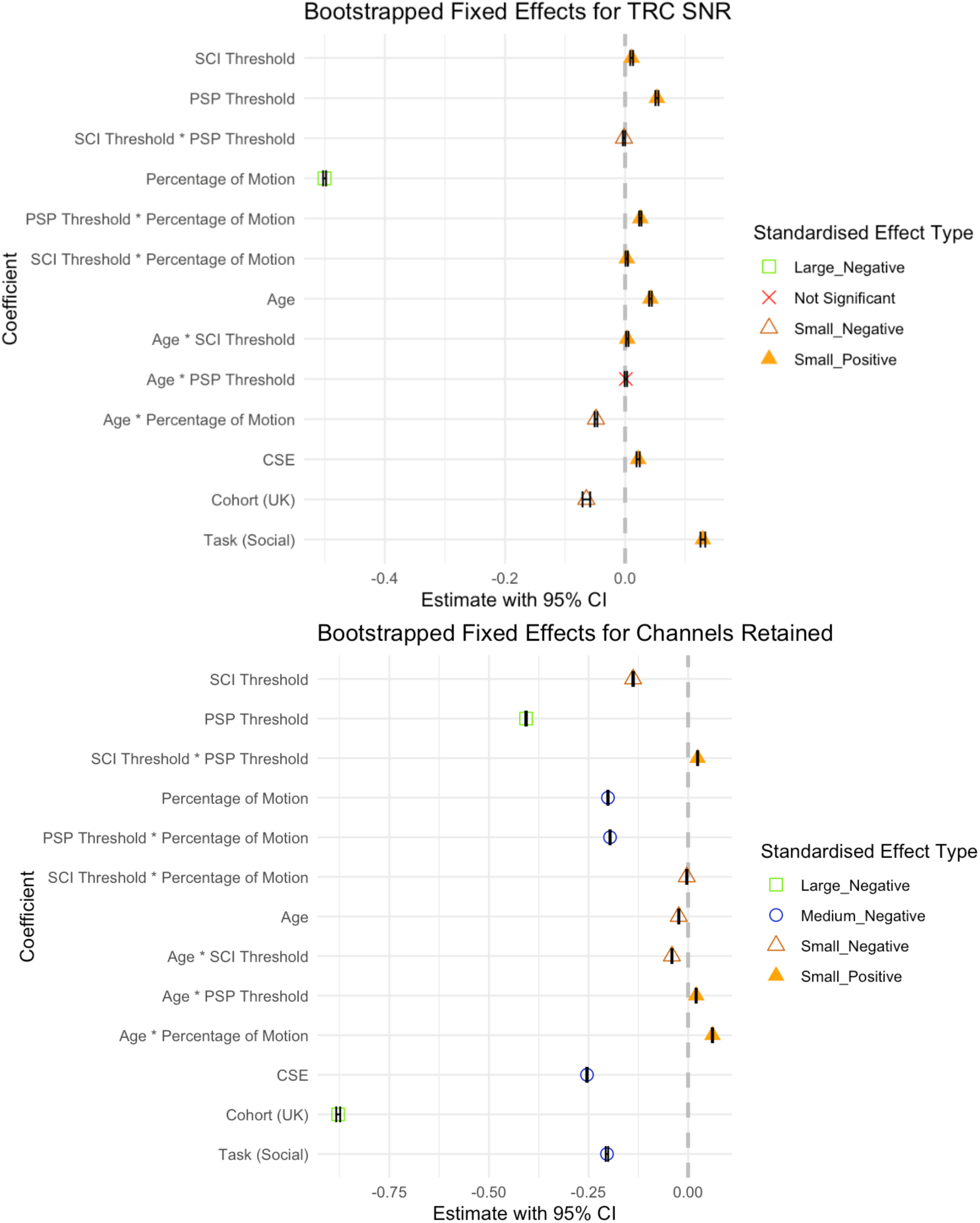
Forest plots showing the results for the effect of each fixed effect in Equation 2 on TRC SNR and Channels Retained, obtained via bootstrapping scaled values. Top: Forest plot showing the type and relative effect size of each fixed effect on TRC SNR. Bottom: Forest plot showing the type and relative effect size of each fixed effect on Channels Retained.

#### 3.2.1 Predictors for TRC SNR

Both SCI Threshold (*β* = 0.0107, 95% CI [0.0084, 0.0131], *SE* = 0.0012) and PSP Threshold (*β* = 0.0530, 95% CI [0.0505, 0.0556], *SE* = 0.0013) had small, positive effects representing an increase in signal quality for higher threshold values. The interaction effect between PSP Threshold and SCI Threshold had a small, negative effect (*β* = −0.0019, 95% CI [−0.0037, −0.0001], *SE* = 0.0009).

PoM had the largest impact on TRC SNR, with a large negative effect (*β* = −0.5004, 95% CI [−0.5028, −0.4979], *SE* = 0.0012) corresponding to a decrease in signal quality in data with more frequent incidences of motion. This decrease in signal quality with motion was offset slightly as participants aged, driven mostly by a more moderate decrease in the 18mo data, illustrated by a small negative interaction effect between and PoM (*β* = −0.0485, 95% CI [−0.0506, −0.4062], *SE* = 0.0011). Small interaction effects between PoM and both SCI Threshold (*β* = 0.0027, 95% CI [0.0008, 0.0047], *SE* = 0.0010) and PSP Threshold (*β* = 0.0253, 95% CI [0.0234, 0.0272], *SE* = 0.0010) exhibited trends which saw high threshold values mitigate the detrimental effect on TRC SNR; the slightly larger main (PSP) and interaction (PSP:PoM) effect in the case of PSP Threshold led to greater mitigation of the TRC SNR decline due to PoM than in the case of SCI (see Figure 5a).

**Figure 5:**
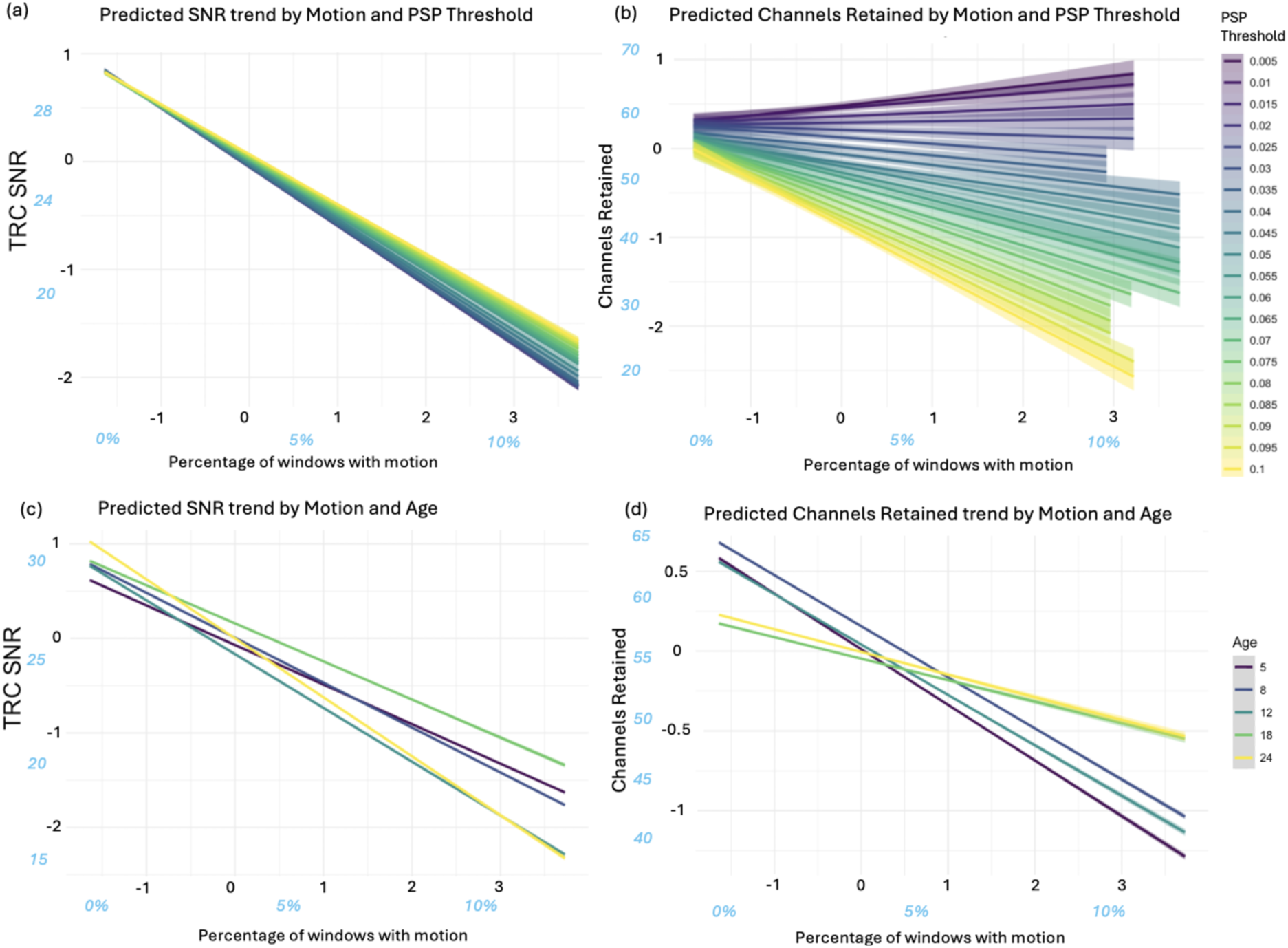
The relationship between motion, and data quality and retention. Blue, italicised axes values represent the approximate original values before scaling during analysis. (a) Predicted TRC SNR trend by Percentage of Motion, grouped by PSP Threshold. (b) Predicted Channels Retained trend by Percentage of Motion, grouped by PSP Threshold. (c) Predicted TRC SNR trend by Percentage of Motion, grouped by Age. (d) Predicted Channels Retained trend by Percentage of Motion, grouped by Age.

The small, negative effect of Cohort (*β* = −0.8783, 95% CI [−0.8832, −0.8733], *SE* = 0.0025) is notable since this fixed effect had larger effects on other outcomes, including Channels Retained. The non-significant effect of PSP:Age (Figure 6b) is notable given the size of the dataset and rarity of non-significant predictors throughout this work.

**Figure 6:**
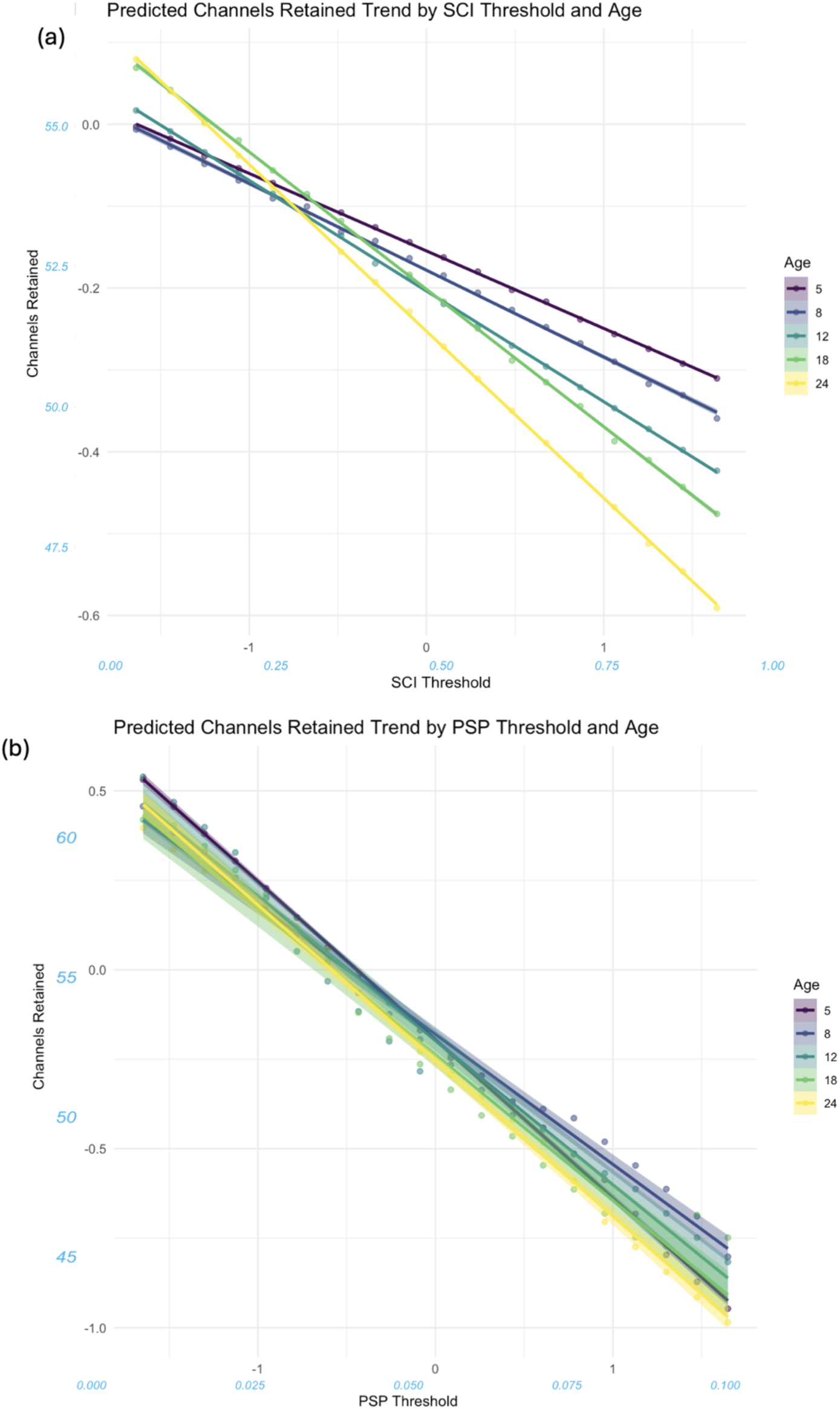
Effect of the interaction between QT-NIRS thresholds and age on channel retention. (a) The interaction between and SCI Threshold, showing the mitigating impact of high SCI values for younger participants. (b) The interaction between and PSP Threshold, in which a pattern with age is harder to ascertain.

#### 3.2.2 Predictors for Channels Retained

Both SCI Threshold (*β* = −0.1381, 95% CI [−0.1401, −0.1361], *SE* = 0.0010) and PSP Threshold (*β* = −0.4067, 95% CI [−0.4087, −0.4048], *SE* = 0.0009) had small and large negative effects on Channels Retained, respectively (see Figure 7). This indicates an increase in the number of channels pruned when using higher threshold values, particularly for the PSP Threshold. The interaction effect between SCI Threshold and PSP Threshold was small and negative (*β* = −0.8783, 95% CI [−0.8832, −0.8733], *SE* = 0.0025), with each threshold alleviating the negative effect of the other at high parameter values.

**Figure 7:**
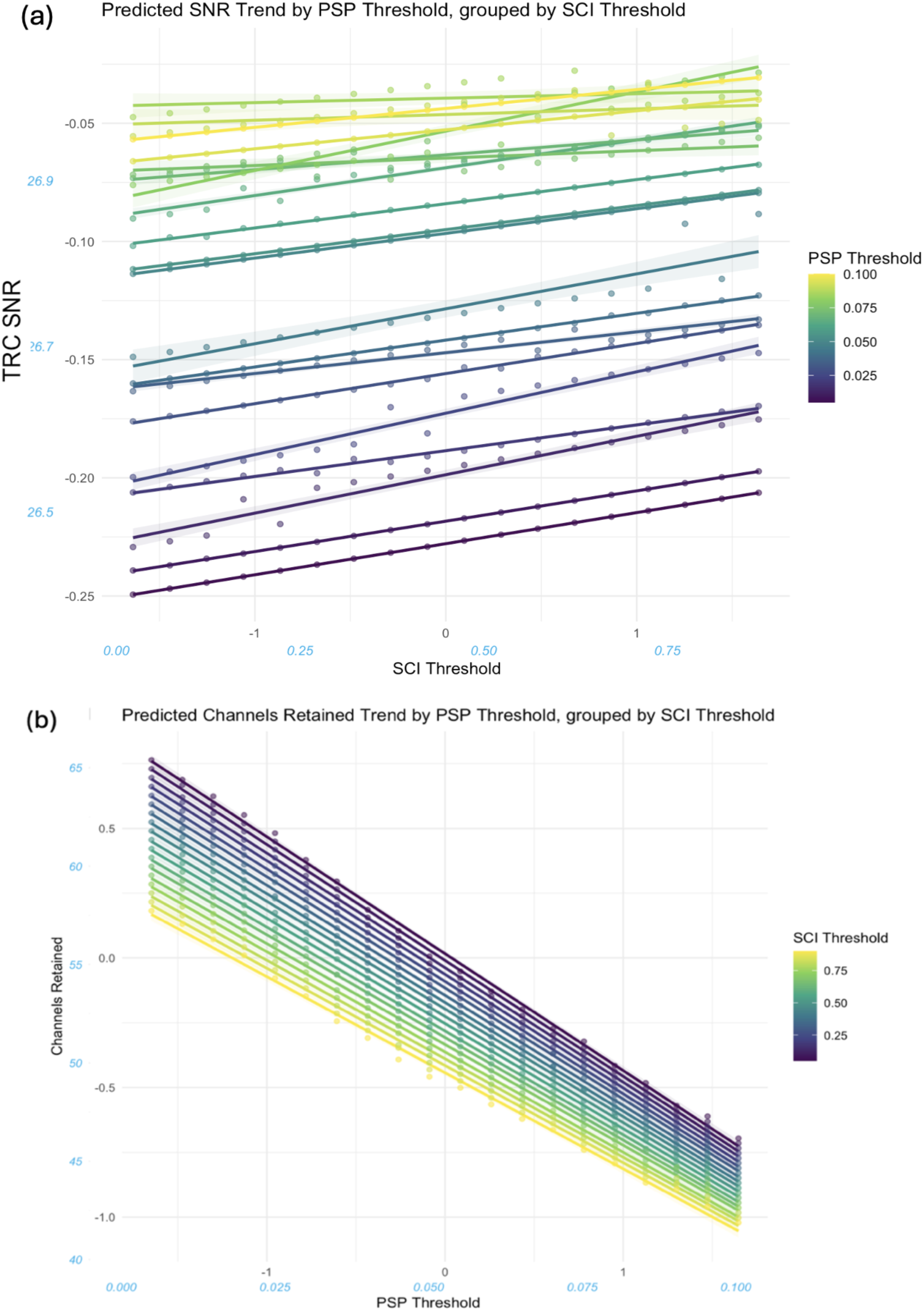
The effect of the interaction between SCI Threshold and PSP Threshold on signal quality and retention. (a) The interaction between thresholds acting on TRC SNR, showing an inconsistent, slight decrease in TRC SNR when compared to the overall trend for high SCI- and PSP Threshold values. (b) The interaction between thresholds acting on Channel Retained, showing a more consistent trend across threshold levels and the dampening of the reduction in channels when both SCI- and PSP Threshold values are high.

PoM had a significant, medium negative impact (*β* = −0.2013, 95% CI [−0.2030, −0.1995], *SE* = 0.0009), with higher amounts of motion associated with decreased channel retention. At the two oldest ages (i.e. 18- and 24mo), greater proportions of motion in the data had a less drastic negative impact on the number of channels pruned (see Figure 5d) which was captured by the small positive interaction effect Age:PoM (*β* = 0.0610, 95% CI [0.0593, 0.6264], *SE* = 0.0009). As with TRC SNR, interaction effects of PoM with both SCI Threshold (*β* = −0.0034, 95% CI [−0.0050, −0.0019], *SE* = 0.0008) and PSP Threshold (*β* = −0.1962, 95% CI [−0.1977, −0.1945], *SE* = 0.0008) moderated this channel reduction for higher percentages of motion. Though both interactions were significant, PSP:PoM was of medium effect size and also acted on two (negative) medium main effects, leading to a near negation of the detrimental impact of motion on predicted Channels Retained for low PSP Threshold values (see Figure 5b).

Task (*β* = −0.2038, 95% CI [−0.2069, −0.2007], *SE* = 0.0015) and Cohort (*β* = −0.8783, 95% CI [−0.8832, −0.8733], *SE* = 0.0025) had medium and large effects on Channels Retained, respectively, resulting in lower channel retention for the SNS task and UK cohort.

## 4 Discussion

Channel pruning in infant fNIRS research is a crucial step in data-analysis, yet often performed with subjective parameters or adult derived thresholds. Moreover, approaches to channel pruning, including QT-NIRS and CV pruning, in infant fNIRS data have often relied on adult-derived thresholds or single-domain measures, without systematic evaluation of their suitability for infant data. The work described here took a quantitative approach to directly compare pruning strategies and parameter settings for infant sparse array data. It was found that QT-NIRS, in its consideration of both time- and frequency-domain signal quality measures, achieved a better balance of data quality and retention than CV pruning. Additionally, parameter values for QT-NIRS were investigated alongside contextual factors, with lower values than those used for adults likely being preferable. These findings supplement work in the wider literature aimed at improving infant NIR imaging data pipelines more broadly [6], [37], [44].

### 4.1 QT-NIRS as a channel pruning method

QT-NIRS produced infant fNIRS data of significantly greater SNR than pruning in all but one age/task/cohort-specific comparison (SNS task in UK infants at 24mo), whilst controlling for data retention. Higher signal quality remained statistically significant when channels were pruned using SCI only, suggesting that evaluating the signal using a method which accounts for a temporal data characteristic (correlation) is more effective than assessment using time-independent measures. It may also indicate that evaluation of the signal in subsampled windows provides a more accurate reflection of signal quality than holistic signal assessment, as was the case with CV pruning. This difference in signal quality, even without using PSP is observed despite evidence suggesting that SCI may be biased by latent, undetected motion in the signal in at least some participants (see Effects of Motion).

Using both SCI- and PSP thresholds further improved signal quality, resulting in higher mean TRC SNR values in all cases. Though non-significant, the higher TRC SNR values reinforce the advantage of incorporating both frequency- and time-domain metrics and provide further justification for the use of QT-NIRS when channel pruning of infant fNIRS data.

### 4.2 Effects of QT-NIRS parameter choice

Higher SCI- and PSP thresholds reduced data retention by pruning more channels, but improved signal quality in TRCs. The effect of parameter changes was smaller than anticipated, particularly on signal quality. PSP Threshold had more influence than SCI Threshold on both outcomes but particularly on Channels Retained, illustrating that caution is needed when altering this threshold to avoid unnecessary data loss.

A positive SCI:PSP interaction reduced the channel pruning rate relative to that expected when considering both thresholds in isolation (see Figure 7b), suggesting that the potential cost to data retention from increasing one threshold value is mitigated when the other value is also high. This aligns with the use of complementary signal metrics in QT-NIRS which must both be of sufficient quality to retain channels: each threshold will exclude channels which may be included by the other, with the overlap in excluded channels increasing with parameter values.

The slight negative SCI:PSP interaction effect on TRC SNR may be driven by the highest values for both parameters (Figure 7a). Higher thresholds increased the likelihood of pruning channels with neuronally-evoked cortical haemodynamic responses and, consequently, high SNR; their removal is likely to reduce the mean TRC SNR more than would be expected for lower threshold values. Change in TRC SNR is also less uniform across SCI- and PSP Threshold values than for Channels Retained. Unlike channel retention, which reflects the whole array, TRC SNR is localized to select channels. Single channels will therefore likely contribute a much greater proportional value to the average TRC SNR and the consequence of pruning these channels is likely to have a larger relative impact on the TRC SNR. Additionally, pruning a channel consistently reduces Channels Retained whereas its impact on TRC SNR depends on the pruned channel’s SNR, further contributing to the inconsistency in change.

### 4.3 Effects of Motion

Substantial negative effects of PoM on both signal quality and channel retention suggest that motion reduces signal quality in at least some channels even when artefacts are not considered in the pruning analysis process, as was the case in this work. Motion artefacts may cause optodes to move, dislodge, or uncouple completely, causing a poorer quality signal reflected in decreased TRC SNR. In turn, channels affected by motion may be pruned to a greater extent during QT-NIRS processing, leading to lower Channels Retained values.

Interaction effects show PSP Threshold had the greater impact on motion-affected data of the two thresholds, improving signal quality but reducing channel retention, especially at high threshold values. In contrast, a comparatively modest effect of SCI Threshold on signal quality and retention was found. Motion exhibited a significant adverse effect on Average PSP, likely due to optode displacement severe enough to disrupt coupling, and a positive effect of motion incidences on Average SCI, suggesting that SCI measures may be capturing correlation induced by latent, undetected motion artefacts in the data. Future work may examine the effect of different motion detection parameters, or alternative motion correction methods altogether such as the Sobel filter [45], acceptance rate adaptive algorithm [46], global variance of temporal derivatives [47], [48] or entropy-based methods [49].

The potential (interaction) effect of high PSP Threshold values on channel retention, especially for channels with a greater proportion of motion, warrants caution for users when looking to employ high PSP Threshold values. This is particularly true given the relatively small beneficial impact on signal quality of increasing the PSP threshold, indicated by its main effect on TRC SNR. This is consistent with the argument for using lower values discussed in the Recommendations section of the Discussion. Conversely, motion incidences have the largest negative impact on TRC SNR; low PSP thresholds may exacerbate this effect if motion is not appropriately addressed.

### 4.4 Effects of age

Age had small effects on TRC SNR and Channels Retained, suggesting limited changes in signal quality and retention in infants between the age of 5 and 24mo. Reflecting this, associated trends were less commonly observed with age than for other predictors; however, it was notable that channel loss was less severe when increasing the SCI Threshold in younger participants (Figure 6). Additionally, while higher signal PoM is associated with reduced channel retention, older infants retained more channels than younger ones for data with high motion prevalence (Figure 5d). This may be due to younger infants manually touching or grabbing the cap more, causing more severe artefacts that permanently displace optodes and affect subsequent coupling, an assertion supported by post-hoc data examination which suggested a decrease in motion severity with age (see Supplementary Materials, Figure S 7)

Interactions with age for TRC SNR showed no meaningful patterns. This may reflect a lack of consistent changes within the age range sampled here. It may also reflect the multifaceted mix of concepts which ‘age’ represents: the interplay of the physiological and behavioural changes with age may be too complex for a simple fixed effect to capture. Interactions between age and other model terms (e.g. CSE, CL) caused convergence issues and were omitted. Future research may prioritize investigating age-related change and associated factors affecting signal quality – such as hair characteristics [23], hair style [31], or hair type changes with age [50].

### 4.5 Other predictors

#### 4.5.1 Cohort

Cohort had substantial effects on signal quality and retention, likely due to factors such as testing environment, tester experience, parent and infant behaviours, and sample size. Since cohort was not a primary predictor, its main effect was not investigated in depth and interaction terms were not included. Nevertheless, data from the Gambian cohort – whose skin and hair characteristics have been found to pose challenges for fNIRS signal quality [23] – exhibited better signal quality and retention, to the extent that data exclusion was far lower for Gambian infants in general and almost non-existent at the infant level (see Figure 2). There may be several reasons for this. Firstly, UK infants generally had finer hair which was longer at later assessment ages, possibly causing cap slippage and poorer subsequent signal quality, as these values appear to decrease with age. Lower motion incidence in Gambian infants may also play a role, possibly due to a relative lack of familiarity with digital screens in daily life leading to an increased focus on a novel, unfamiliar object. While physical characteristics of participants undoubtedly affect the fNIRS signal, cohort differences in this study emphasize the need to consider other factors which affect signal quality during data collection.

#### 4.5.2 Weak or saturated channel signals

A small and significant negative effect of CSE on channel retention was anticipated, considering it is itself a measure of channel removal. CSE was also significantly negatively associated with data quality, however, suggesting poor coupling in channels with extremely low quality signal could be affected by – or causing – signal quality issues elsewhere in the array. It may be of interest to investigate whether signal quality was worse in channels located closely to the CSE channels in future work, as has been the focus of prior work into motion artefact detection [49].

### 4.6 Strengths and limitations

A key strength of this work is the dataset used. Data encompassed five testing time points across the age range 5- to 24mo, allowing assessment of QT-NIRS and its key parameters for infants whilst accounting for age-related structural changes in skull thickness, cardiac signal properties, and surface vasculature [51]. Data from two sites were also assessed, one of which was a rural setting in a sub-Saharan African country, addressing a common bias which often exists in infant neuroimaging studies when participant recruitment is limited to predominantly White infants from high-income contexts [52]. As such, the findings are likely to be more robust and generalisable than those derived from smaller or single-site samples. The inclusion of longitudinal data allows for the assessment of within- and between-participant changes over time, providing richer insights into the effect of processing methods than would be possible from cross-sectional analyses alone.

Another strength is the MLM approach which accounted for the hierarchical variance structure of longitudinal data grouped by task and cohort, reducing bias and enabling random intercepts and slopes during regression analysis to capture individual differences [39]. This approach also handled missing data introduced through prior channel removal (CSE channels), missed visits or incomplete testing, which would not have been possible with many other common analysis procedures [53]. Domain knowledge and data-driven insight were combined to prioritize model predictors and validate model assumptions and additionally fitted and assessed subsidiary MLMs to ensure the final model (Equation 2) was as comprehensive as possible.

QT-NIRS was compared with CV pruning, using a maximum wavelength CV difference of 0.2, as previously applied with data from this study [14]. Significant differences in signal quality between QT-NIRS and CV pruning were found in all but one age/task/cohort combination. However, another common CV pruning approach uses a single-channel threshold instead, pruning both channel wavelength signals at least one of them exceeds it [12], [32], [54]. Future work could investigate whether the significant differences found here persist when using this alternative thresholding method.

The focus of this work was on optimizing thresholds of the two QT-NIRS parameters which are most pivotal to performance, for which only reported recommended values for adult participants were found in the literature. Future work may focus on other parameters, such as changing the quality threshold (*q_threshold*), balancing channel quality discernment with the risk of losing nuance in signal characteristics. Future studies could also investigate the impact of altering window size or overlap: the former will likely balance measurement accuracy within windows against overall temporal sensitivity; the latter may provide more window temporal sensitivity at the expense of computational efficiency. Given the dominance of PSP Threshold change on outcomes reported here, it may also be of interest to explore pruning using only the PSP threshold [55].

This work must also be placed in the context of the increasing impact of machine learning on infant NIR imaging data processing, with future work in the field likely to assess the efficacy and generalisability of such methods. Deep learning approaches are frequently being added to the literature: a machine learning based detector has been developed to identify ‘bad’ channels to be pruned, for example [55]. This approach was shown to be more adaptive, interpretable, and effective across diverse noise types than QT-NIRS and other methods, so future work may seek to assess the efficacy of this independently. In mitigation, care should be taken to ensure limitations common to many deep learning approaches (e.g. overfitting) are addressed through appropriate validation strategies, including the use of independent test sets and cross-validation.

Model design was limited by convergence issues, particularly when including random slopes and intercepts. The most likely and important sources of variation were prioritized, however it was not possible to fully include all relevant interaction terms or random effects. Additionally, linear terms were used in the models to simplify interpretability. Future work may incorporate non-linear terms, as an alternative to the bootstrapping approach for dealing with non-normal residuals. Other alternative approaches could include sensitivity analysis of the influential points and outliers; variance modelling for specific predictors; and utilisation of robust standard errors.

While age is likely to reflect more than behaviour and motion, the longitudinal study design may also result in participant familiarity with the testing procedures and the paradigms, in turn possibly affecting behaviour, attention, stress levels, engagement, and – consequently – the recorded data. Thus, caution should be taken when examining the effects of the age model term and its interactions, recognising this as a potential confounding factor in this work.

Further evaluation of QT-NIRS as a channel pruning method for infant fNIRS data is still necessary to address the limitations of this work, extend it to high-density systems, and compare it to alternative approaches including those incorporating machine learning [55].

### 4.7 Recommendations

Based on this work, the following guidelines for channel pruning infant fNIRS data are recommended:

(1) QT-NIRS as a pruning method is preferable to pruning using CV (when using a minimum threshold difference between wavelengths)
(2) Users should conduct channel pruning on motion-free data, with considerable emphasis during initial preprocessing placed on adequately identifying motion artefacts
(3) Tuning of the PSP Threshold should be prioritized over the SCI threshold in data
(4) To provide a good trade-off between data quality and retention, lower thresholds (with minima of *psp_threshold* ≈ 0.04–0.05 and *sci_threshold* ≈ 0.6) can be used for infant data than those recommended for adults, especially in older infants with fine/slippery hair, since (amongst other factors):

(i) There are a large proportion of data showing SCI- and PSP values lower than the adult thresholds of 0.8 and 0.1, respectively (Figure 2)
(ii) The risk of removing data with higher thresholds is likely greater than the potential gain in improved signal quality
(iii) There is a plateau in both average TRC SNR and Channels Retained values when using lower threshold values than the advised minima (see Supplementary Materials, Figure S 6)

A pictorial guide to the effect of the most common predictors on data quality and retention is also included in Figure 8.

**Figure 8:**
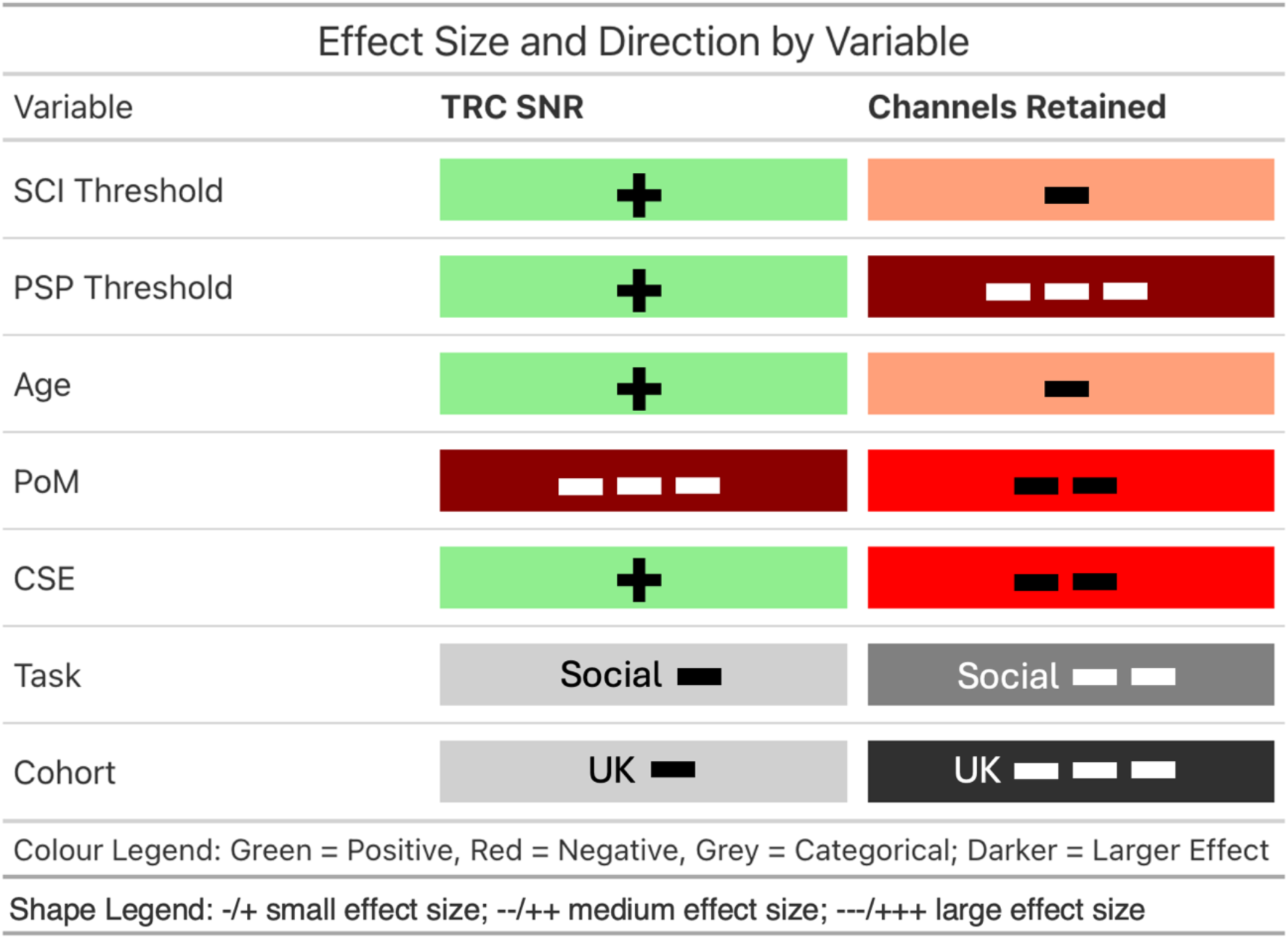
Guide to the effects of the parameters and data characteristics on data quality and retention. First five rows represent positive (green) or negative(red) associations with increasing numerical parameters (SCI Threshold and PSP Threshold) or data characteristics (Age, PoM, CSE). All positive effects are small in size. Bottom two rows represent categorical variables, with greyscale shading indicative of the impact on outcome changing the categorical variable may have.

It is still strongly recommended that fNIRS users take the appropriate time to understand the dataset being analysed since no approach can possibly be universal. To aid final parameter selection, a tool which other users may find useful to guide parameter selection was developed. Alongside code used during processing and analysis reported in this study can be found at: https://github.com/sam-beaton/pruningComparisons/. The tool is designed to examine the effects of parameter threshold changes within a group (e.g. age), by establishing a trend capturing the trade-off between data quality and retention, then assessing which specific SCI- and PSP Threshold parameter combinations perform best in relation to this trend.

## 5 Conclusion

Recent advances in fNIRS channel pruning approaches show promise for improving the accuracy of preprocessing by evaluating both the time and frequency domains. The work described here compared QT-NIRS with an established pruning method utilising CV, across 5 infant ages, two paradigms, and two sites. It was found that QT-NIRS provides data with greater signal quality when controlling for data retention.

The consequences of different parameter choices for QT-NIRS were also demonstrated, highlighting the importance of the PSP Threshold, plus the influence of motion. Evidence-based recommendations for QT-NIRS pruning, and parameter choice for infant data with different characteristics, are provided.

## Code and Data Availability

All data used in the analyses presented can be made available following relevant approvals. The code used to conduct the analyses and generate the figures presented in this paper are available at https://github.com/sam-beaton/pruningComparisons. Code relies on the following open source R libraries: arules [56], broom [57], car [58], data [59], doParallel [60], dplyr [61], effectsize [62], effects [58], e1071 [63], foreach [64], ggplot2 [65], ggpubr [66], gridExtra [67], lme4 [41], lmerTest [68], MASS [69], matrix [70], MuMIn [71], Scales [72], tidyr [73], viridis [74].

## Acknowledgments

We would like to place on record our thanks to the children, mothers and wider families who took part in this study. In addition, we would like to thank the data collection teams in both The Gambia and the UK. Finally, we thank Luca Pollonini for his contribution to early discussions concerning the framing of the work, and Johann Benerradi, for providing task-relevant channels for the social/non-social task prior to their publication.

## Author Contributions (CRediT taxonomy)

SB: conceptualisation, formal analysis, methodology, software, validation, visualisation, writing – original draft, writing – review and editing; BB: conceptualisation, methodology, supervision, writing – review and editing; CB: conceptualisation, methodology, writing – review and editing; CE: funding acquisition, project administration, resources, writing – review and editing; SLF: funding acquisition, project administration, resources, writing – review and editing; EM: data curation, investigation, project administration; SMc: data curation, investigation, project administration, supervision; ABR: conceptualisation, data curation, supervision, visualisation, writing – review and editing; SM: funding acquisition, project administration, resources, supervision, writing – review and editing.

## Biographies

**Samuel Beaton** is a doctoral researcher and research assistant at King’s College London, developing analytical pipelines for functional near-infrared spectroscopy (fNIRS) and high-density diffuse optical tomography (HD-DOT) data.

**Chiara Bulgarelli** is a Senior Lecturer at the Centre for Brain and Cognitive Development at Birkbeck, University of London. She has over a decade of experience using fNIRS with infants and toddlers to study the neural mechanisms underlying social interactions and the development of brain connectivity. Recently, she pioneered the use of fNIRS in non-traditional lab settings, such as with toddlers interacting in a virtual reality environment.

**Borja Blanco** is a postdoctoral research associate in the Department of Psychology at the University of Cambridge. His work focuses on developing data processing and analysis methods for optical neuroimaging in developmental populations. He applies these methods to investigate infant functional brain development and the contextual factors that influence it.

**Clare Elwell** is a professor of medical physics leading projects to investigate brain function using functional near infrared spectroscopy in high and low resource settings in adult and infants.

**Sarah Lloyd-Fox** is a Principal Research Associate in the Department of Psychology, University of Cambridge. She leads several multi-disciplinary projects focusing on developmental trajectories of early cognitive and brain development during pregnancy, infancy and early childhood. Her research focuses on understanding how family and environmental context - i.e. contextual factors such as poverty associated challenges and enriched multigenerational family support - shape early life.

**Ebrima Mbye** is a Field Coordinator at MRCG@LSHTM, formerly employed on the BRIGHT Project and currently engaged on the INDiGO Trial working under Professor Sophie Moore.

**Samantha McCann** is a Public Health Registrar and formerly Postdoctoral Research Associate in the Department of Women and Children’s Health at King’s College London. Her main research interest is supporting early child development, with a strong focus on the impact of undernutrition in infancy on long-term neurodevelopmental outcomes.

**Anna Blasi** is a Postdoctoral Research Fellow at UCL. Her research interests are centered on functional aspects of human physiology. Her research career started with models of the cardiovascular system and the effects of disease. Through her work at UCL, KCL, and Birkbeck, her research interests have shifted toward the use of functional imaging (fNIRS, fMRI) to study brain function and neurocognitive development in early infancy.

**Sophie Moore** is Professor of Global Women and Children’s Health in the Department of Women & Children’s Health at King’s College London and an Honorary Associate Professor at the London School of Hygiene and Tropical Medicine (LSHTM). Her research focuses on the nutritional regulation of ‘healthy’ fetal and infant growth, incorporating infant immune and brain development as outcomes, and on the mechanisms through which maternal, infant and childhood nutrition may influence development and later health.

