## Supplementary Materials for "Investigating the effect of channel pruning on functional near-infrared spectroscopy data collected from children aged 5-24 months"

### Supplementary Materials 1: Descriptions of, and Rationale for, MLM models

#### Supplementary Materials 1.1: Measures, Outcomes and Effects

Considering the anticipated data quality/inclusion trade-off which is central to decision making around preprocessing of fNIRS data (Gemignani and Gervain, 2021), we assessed the performance of pruning methods and parameters using two metrics: (i) signal-to-noise ratio (SNR) and (ii) channel inclusion/exclusion.

We were additionally interested in the effects of other factors commonly relevant for those working with infant participants, such as age, motion, and signal extrema. As such, particular attention was paid to their influence as fixed effects on signal quality and retention during the investigation into the effect of QT-NIRS parameter choices, building on the subsidiary investigations into motion and average coupling metrics. Other measures were more particular to our dataset - such as task, cohort and optode position. We included them to account for the variability they may cause instead.

Indicators contained in brackets for each measure in the following measure descriptions indicate whether they were used as outcome variables (O), predictors (P), or covariates (C). Those with multiple indicators are utilized for different purposes depending on the aims of the assessment in question. We start by describing channel-level measures as these are subsequently built on to inform some of the participant-level measures which follow. Throughout, we worked only with artefact-free data.

#### Supplementary Materials 1.2: Channel-level measures

##### Supplementary Materials 1.2.1: Average SCI and PSP per channel (O)

Average SCI was calculated, for every non-CSE channel, by taking the mean of SCI values across all included time windows. Average PSP was calculated in the same way as Average SCI, using PSP instead of SCI values per time window.

##### Supplementary Materials 1.2.3: Percentage of motion (O), (P)

Motion detection was conducted on each non-CSE channel so that relevant calculations could be performed on motion-free data. We stored the number of windows in which motion was detected as a percentage of the total number of windows for each channel, providing a measure of the prevalence of motion which we named 'Percentage of Motion' (PoM).

##### Supplementary Materials 1.2.4: Channel Location (C)

Posterior channels of the array are located on areas of the skull with more pronounced curvature whereas those located frontally are more likely to be in contact with hair-free areas of scalp in the forehead, likely decreasing and improving coupling, respectively. Variation in signal may also occur due to differences in propagation of the cardiac signal across the head from the rear to the front. To account for this variability we assigned a number from 1-11 to each channel depending on their anterior-posterior location, with lower values representing channels located posteriorly: channels 18 and 37, for example, were assigned ‘1’; channels 15, 19, 34 and 38 were assigned ‘2’, and so on (higher/lower values have no inherent meaning other than to provide a range representing anterior-posterior location). This was labelled ‘Channel Location’ (CL).

**Supplementary Materials 1.2.5: Task-relevant channel signal-to-noise ratio (O)**

Signal quality after pruning was measured using the signal-to-noise ratio in task-relevant channels (TRC SNR). We use the following definition of SNR:

$$SNR = 20\frac{\mu}{\sigma}$$

where $\mu$ and $\sigma$ are the mean and standard deviation of the signal, respectively. We calculated SNR only in ‘task-relevant channels’ (TRCs) – the name we give to channels where we would expect to see a true haemodynamic signal, for brevity. Age-specific TRCs were determined using prior analyses of the two paradigms for both cohorts: the SNS channels were taken from work by Benerradi and colleagues [26] whereas the HaND channels were taken from work by Blasi and colleagues [26]. In line with increased calls in the fNIRS community for consideration of both oxy- and deoxyhaemoglobin (HbO and HbR) chromophores during analysis (Tachtsidis and Scholkmann, 2016), channels were included in the set of TRCs for each age and task if they exhibited a significant evoked haemodynamic response in both HbO and HbR chromophores. The exception to this was the SNS task at 18mo as only 1 channel met this criterion; in order to utilize a comparable number of channels for each age and avoid the inability to assess signal quality should this single channel be pruned, we added channels to the TRC set for this age if activation was exhibited for at least two other ages for the same task.

**Supplementary Materials 1.2.6: Channels retained (O), (P)**

We summed the number of channels per participant included after pruning using the described method and parameter(s) – this was labelled ‘Channels Retained’ (CR).

**Supplementary Materials 1.3: Participant-level measures**

**Supplementary Materials 1.3.1: Age (P)**

Age was included as a five-level predictor (5-, 8-, 12-, 18-, 24mo) to assess change with age.

**Supplementary Materials 1.3.2: Cohort (P)**

Data from two cohorts are incorporated into the analysis in this work. We examined the effect of cohort on optode coupling, mostly due to the potential impact of differing participant physical characteristics and behaviour on signal quality (see 5.1.2 and 5.1.3), so we included cohort as a two-level predictor (GM and UK).

**Supplementary Materials 1.3.3: Task (C)**

Two paradigms included are performed as part of the BRIGHT protocol and included in this work. We are not specifically interested in the effects of the paradigms themselves, therefore we included task as a two-level covariate (HaND and SNS) to account for the contribution to variance of the different paradigms.

#### Supplementary Materials 1.3.4: SCI Threshold (P)

For each run of pruning with QT-NIRS, we used a different *sci_threshold* value, ranging from 0.05 to 0.9, with increments of 0.05. We selected an upper threshold of 0.9 as it represents a value between the recommended value for adult participants (0.8) and the theoretical maximum of SCI = 1, which is considered unrealistic. Initial exploratory analyses for each age/task/cohort combination (see Supplementary Materials 3) – in which different parameter choices were utilized without prior channel extrema pruning – indicated that even very low SCI values continued to alter signal quality, so we opted to use the entire range of lower values.

#### Supplementary Materials 1.3.5: PSP Threshold (P)

We varied the *psp_threshold* value, ranging from 0.005 to 0.1, with increments of 0.005. We used an upper value of 0.1 as it is the upper value recommended for adult participants. We used values which spanned the entire possible range lower than this based on initial results from simpler MLM analyses and the considerable proportion of infant PSP values lower than this value.

#### Supplementary Materials 1.3.6: Channels pruned due to signal extrema (C)

As a preliminary step for both methods, pruning based on minimum recorded values was conducted by discarding any channel in which the minimum light intensity value in the channel was less than 3e-4. This value is informed by previous experience with the NTS system and a subset of the same HaND task data used in the current study [14]. We label these 'channels with signal extrema' (CSE) for brevity; the number of CSE per participant was stored as a participant-level covariate and proxy measure of particularly poor optode coupling.

#### Supplementary Materials 1.3.7: Average Motion Percentage per participant (P), (O)

When utilized in the final MLM assessment to gauge the impact of *sci_threshold* and *psp_threshold* on signal quality and retention with age (see 5.5.1.4), the mean across channels of average PoM windows per channel was used as predictors as this analysis was conducted at the participant-level

### Supplementary Materials 2: Rational for SCI and PSP threshold values

The impact of threshold values on mean TRC SNR values across participants, separated by cohort/age/paradigm combination. Colour bar in each case represents TRC SNR values yielded by each combination of *sci_threshold* and *psp_threshold*.

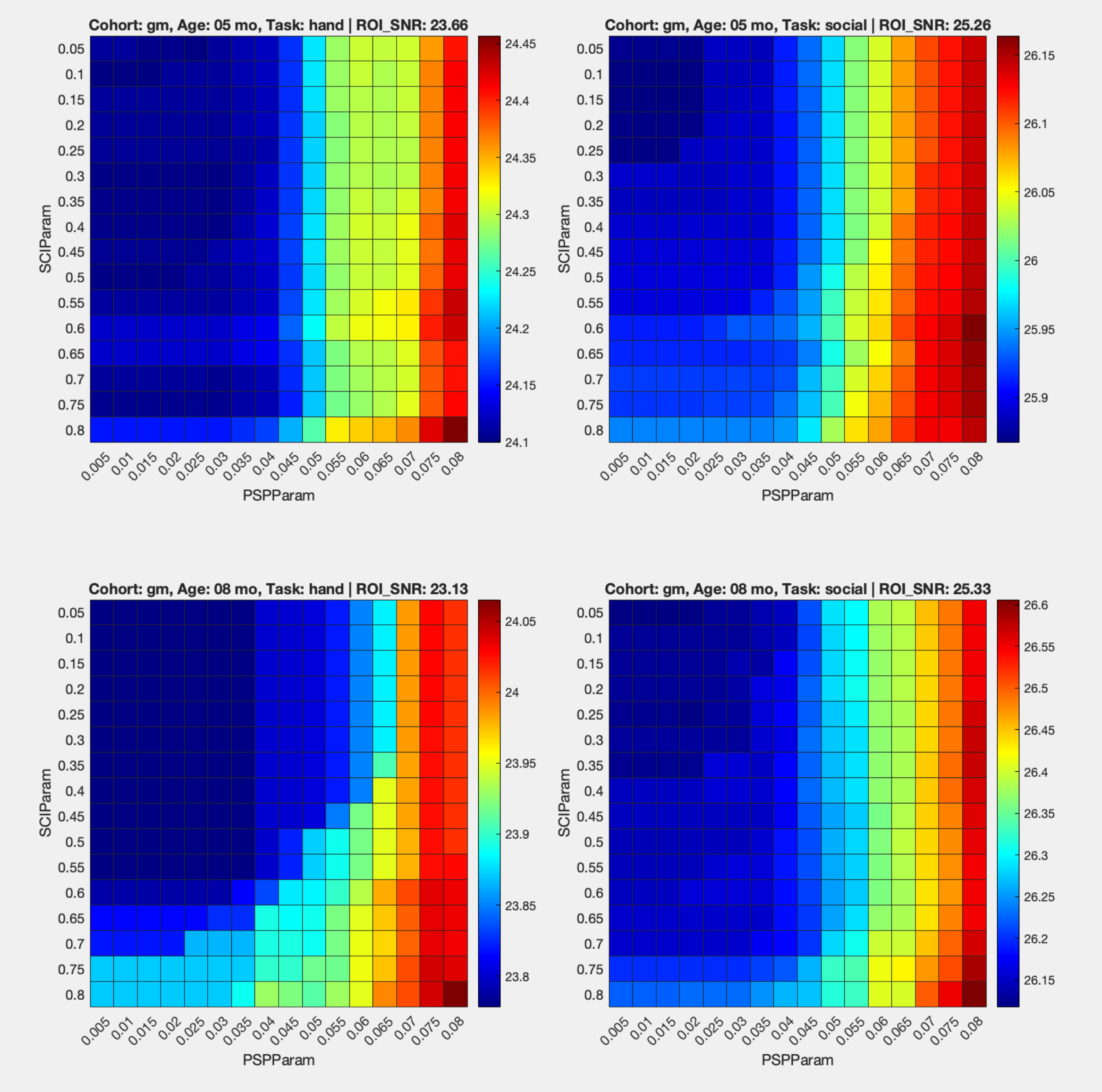


*Figure S 1*

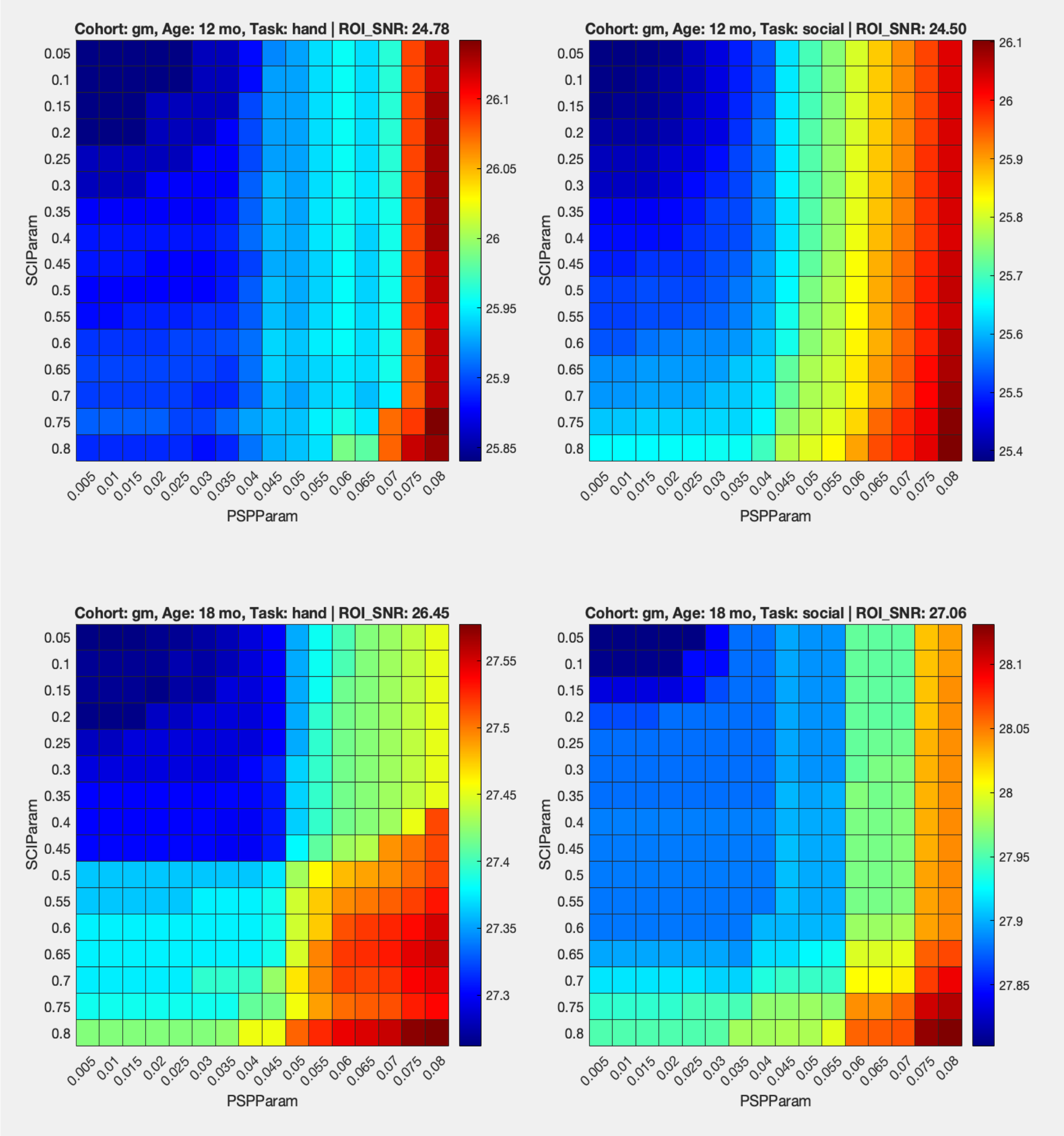
Cohort: gm, Age: 12 mo, Task: hand | ROI_SNR: 24.78
Cohort: gm, Age: 12 mo, Task: social | ROI_SNR: 24.50
Cohort: gm, Age: 18 mo, Task: hand | ROI_SNR: 26.45
Cohort: gm, Age: 18 mo, Task: social | ROI_SNR: 27.06
SCIParam
PSPParam

*Figure S 2*

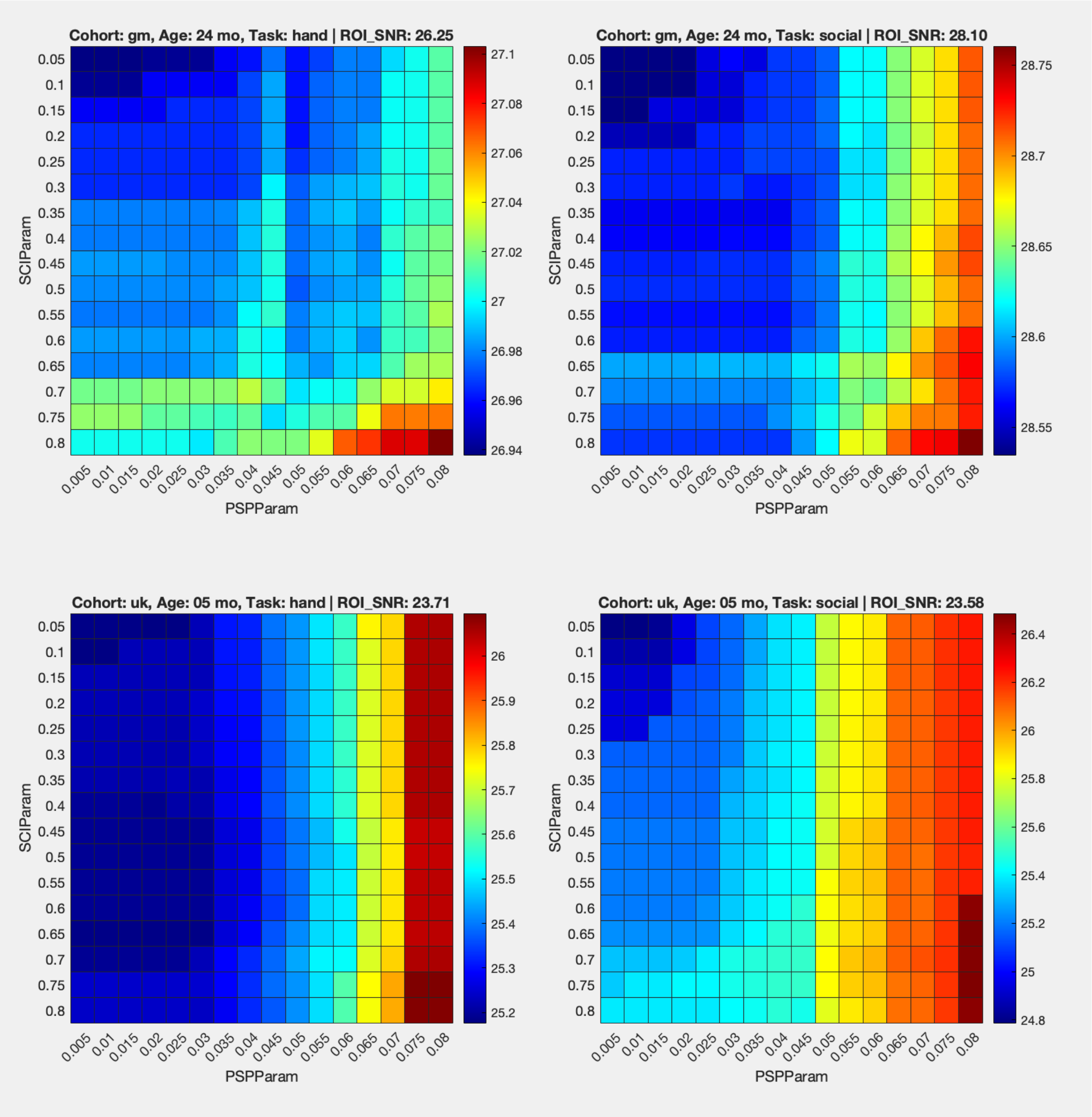
Cohort: gm, Age: 24 mo, Task: hand | ROI_SNR: 26.25
Cohort: gm, Age: 24 mo, Task: social | ROI_SNR: 28.10
Cohort: uk, Age: 05 mo, Task: hand | ROI_SNR: 23.71
Cohort: uk, Age: 05 mo, Task: social | ROI_SNR: 23.58
SCIParam
PSPParam

*Figure S 3*

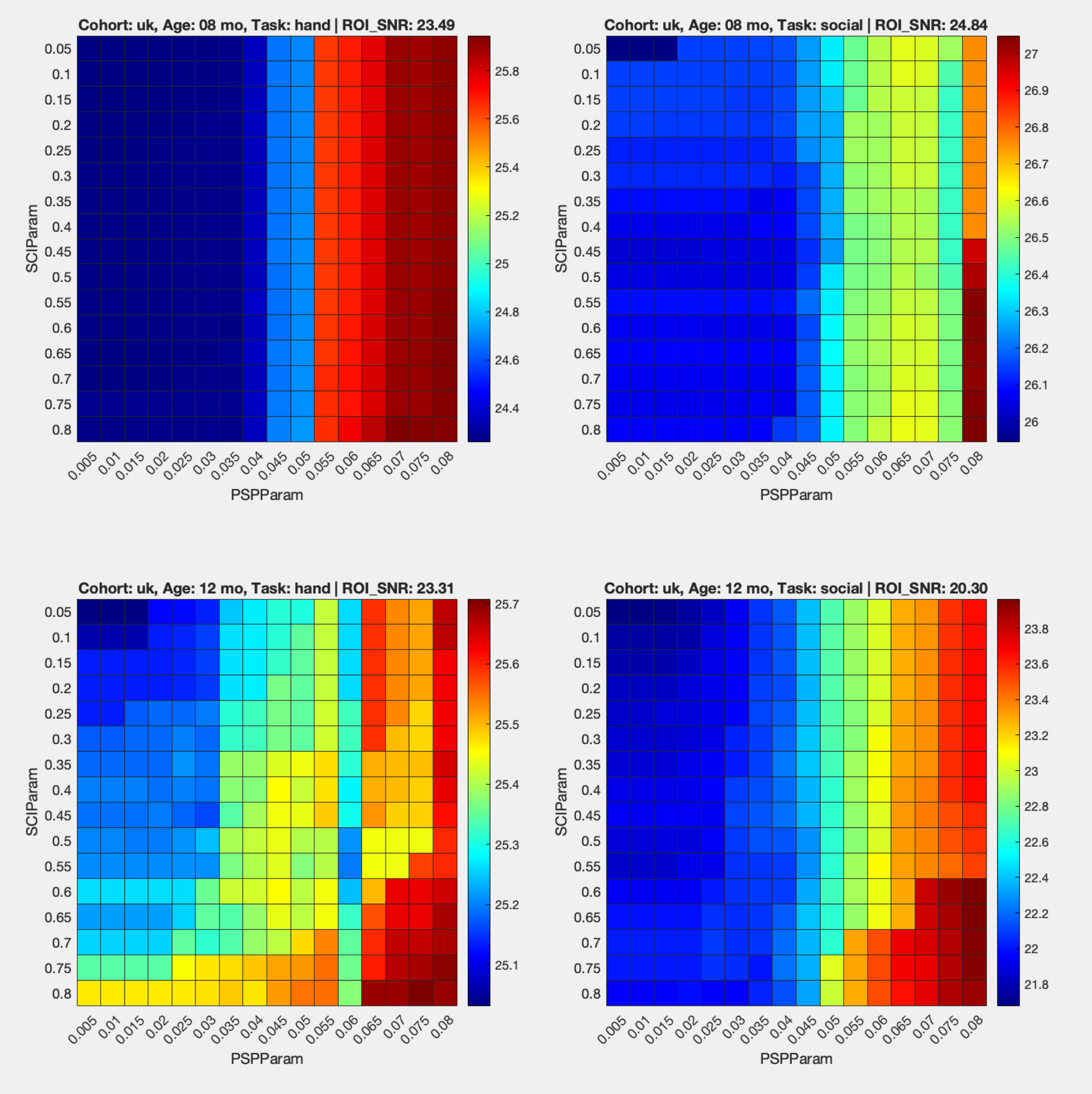


*Figure S 4*

### Supplementary Materials 3: Subsidiary models and analyses

#### Supplementary Materials 3.1: Effect of age on motion

Objective (b)(i) was in part set out to better understand the influences on motion and, therefore, allow us to make more informed choices when modelling the effect of QT-NIRS parameter choices over time.

To address this, we examined the effect of age, task, cohort and channel orientation. As well as the change in behaviour by participant age, we also anticipated that the task may influence participants' engagement due not only to the stimuli involved (the HaND is an auditory task, while the SNS has both visual and auditory stimuli) but also because the HaND paradigm was always presented after the SNS paradigm as part of an extensive battery of assessments. If influential, we supposed that the effect of the task may change with age and repeated data collection sessions as the participants became more familiar and, perhaps, used to the stimuli. We also anticipated that participants in rural Gambia would have been less exposed to digital screens and the other equipment used during data collection than the UK ones , which may affect engagement. This may also change with age, as infants possibly become more familiar over time with the equipment used due to repeated testing and increased exposure in daily life. Finally, we anticipated that channels with optodes in easier to reach positions may influence motion; we also included an interaction term with age for this predictor but model fit was better without it. Our final model to examine the effects on the percentage of motion (PoM) using channel-level data was:

$$Percentage\ of\ Motion \sim Age * Task + Age * Cohort + Optode\ Orientation + (1 \mid ID)$$

*Equation 4*

for data scaled using Equation 3. The generated bootstrapped results were based on 5000 datasets constructed via sampling with replacement of the entirety of the data.

### Supplementary Materials 3.2: Effect of age on average SCI and PSP

We also wanted to understand the predictors for Average SCI and PSP measures, in line with objective (b)(ii). This would also provide further insight into range of values likely to be utilized for *sci_threshold* and *psp_threshold* (objectives (b)(iii) and (c)) by considering the outcome variables of Average SCI and Average PSP themselves, plus demonstrate the effect of motion – if any – on these signal measures (objective (b)(i)).

We included similar predictors and covariates to those used to investigate predictors for motion, namely Age, Task, Cohort, and Channel orientation. We included CSE as a covariate, since we believed it was unlikely to be the case that extremely poor coupling in other optodes would occur in isolation. We also included Percentage of Motion and its interaction with Age as predictors due to the outcome of the investigation into motion. Since we believed motion would influence SCI and PSP measures as discussed in the Introduction, for similar reasons to those used to justify their inclusion in the investigation into effects on motion, we included interaction terms with age for task, cohort and channel location. The final models, for channel-level data, after model fit assessment were:

$$Average_{SCI} \sim Age * Cohort + Age * Task + Age * Percentage\ of\ motion + Age * Optode\ Orientation + CSE + (1 \mid ID)$$

*Equation 5*

$$Average_{PSP} \sim Age * Cohort + Age * Task + Age * Percentage\ of\ Motion + Age * Optode\ Orientation + CSE + (1 \mid ID)$$

*Equation 6*

for data scaled using Equation 3. The generated bootstrapped results were based on 5000 datasets constructed via sampling with replacement of the entirety of the data.

### Supplementary Materials 4: Full results of CV, QT-NIRS (SCI Only) and QT-NIRS pruning comparisons

TRC SNR obtained using each pruning method outlined (CV pruning, QT-NIRS using both parameters, QT-NIRS using *sci_threshold* only).

| Age | Cohort | Task | Method | SCI_Param | PSP_Param | Mean | IQR | SD |
|---|---|---|---|---|---|---|---|---|
| 5 | Gambia | HaND | CV | - | - | 22.066 | 4.647 | 3.303 |
| 5 | Gambia | HaND | SCI | 0.8 | - | 24.117 | 5.244 | 3.482 |
| 5 | Gambia | HaND | QT | 0.2 | 0.04 | 24.271 | 5.095 | 3.484 |
| 8 | Gambia | HaND | CV | - | - | 21.599 | 4.839 | 4.438 |
| 8 | Gambia | HaND | SCI | 0.9 | - | 23.805 | 5.716 | 4.729 |
| 8 | Gambia | HaND | QT | 0.8 | 0.05 | 23.922 | 5.689 | 4.700 |
| 12 | Gambia | HaND | CV | - | - | 22.937 | 4.540 | 4.115 |
| 12 | Gambia | HaND | SCI | 0.85 | - | 25.878 | 4.640 | 4.009 |
| 12 | Gambia | HaND | QT | 0.05 | 0.055 | 25.987 | 4.233 | 3.947 |
| 18 | Gambia | HaND | CV | - | - | 24.696 | 5.190 | 4.094 |
| 18 | Gambia | HaND | SCI | 0.1 | - | 27.342 | 5.117 | 3.912 |
| 18 | Gambia | HaND | QT | 0.1 | 0.025 | 27.428 | 4.884 | 3.876 |
| 24 | Gambia | HaND | CV | - | - | 24.580 | 5.900 | 4.622 |
| 24 | Gambia | HaND | SCI | 0.75 | - | 26.992 | 5.107 | 4.518 |
| 24 | Gambia | HaND | QT | 0.35 | 0.05 | 27.020 | 4.904 | 4.501 |
| 5 | Gambia | Social | CV | - | - | 23.771 | 4.502 | 3.958 |
| 5 | Gambia | Social | SCI | 0.55 | - | 25.900 | 5.285 | 3.852 |
| 5 | Gambia | Social | QT | 0.4 | 0.035 | 26.027 | 5.300 | 3.788 |
| 8 | Gambia | Social | CV | - | - | 23.928 | 5.623 | 4.029 |
| 8 | Gambia | Social | SCI | 0.55 | - | 26.165 | 6.165 | 4.261 |
| 8 | Gambia | Social | QT | 0.55 | 0.02 | 26.363 | 6.061 | 4.163 |
| 12 | Gambia | Social | CV | - | - | 22.930 | 5.635 | 4.226 |
| 12 | Gambia | Social | SCI | 0.6 | - | 25.511 | 6.161 | 4.515 |
| 12 | Gambia | Social | QT | 0.15 | 0.045 | 25.803 | 5.717 | 4.408 |
| 18 | Gambia | Social | CV | - | - | 25.535 | 6.068 | 4.365 |
| 18 | Gambia | Social | SCI | 0.25 | - | 27.896 | 5.399 | 4.202 |
| 18 | Gambia | Social | QT | 0.1 | 0.035 | 27.979 | 5.383 | 4.173 |
| 24 | Gambia | Social | CV | - | - | 26.829 | 5.907 | 4.367 |
| 24 | Gambia | Social | SCI | 0.05 | - | 28.575 | 4.704 | 4.250 |
| 24 | Gambia | Social | QT | 0.05 | 0.005 | 28.647 | 4.774 | 4.238 |
| 5 | UK | HaND | CV | - | - | 22.440 | 3.816 | 4.505 |
| 5 | UK | HaND | SCI | 0.05 | - | 25.214 | 3.827 | 3.240 |
| 5 | UK | HaND | QT | 0.05 | 0.005 | 25.605 | 3.510 | 3.243 |
| 8 | UK | HaND | CV | - | - | 22.458 | 4.719 | 4.211 |
| 8 | UK | HaND | SCI | 0.85 | - | 24.277 | 6.761 | 5.088 |
| 8 | UK | HaND | QT | 0.85 | 0.03 | 25.120 | 5.856 | 4.519 |
| 12 | UK | HaND | CV | - | - | 22.265 | 3.617 | 5.594 |
| 12 | UK | HaND | SCI | 0.4 | - | 25.234 | 4.937 | 4.275 |
| 12 | UK | HaND | QT | 0.2 | 0.03 | 25.445 | 4.742 | 4.206 |
| 18 | UK | HaND | CV | - | - | 24.565 | 4.743 | 3.307 |
| 18 | UK | HaND | SCI | 0.05 | - | 26.918 | 6.058 | 3.823 |
| 18 | UK | HaND | QT | 0.05 | 0.005 | 27.245 | 5.732 | 3.813 |
| 24 | UK | HaND | CV | - | - | 25.869 | 4.431 | 5.996 |
| 24 | UK | HaND | SCI | 0.05 | - | 28.138 | 4.380 | 4.861 |
| 24 | UK | HaND | QT | 0.05 | 0.005 | 28.349 | 4.088 | 4.765 |
| 5 | UK | Social | CV | - | - | 22.924 | 4.773 | 5.192 |
| 5 | UK | Social | SCI | 0.25 | - | 25.151 | 4.126 | 4.681 |
| 5 | UK | Social | QT | 0.35 | 0.005 | 25.815 | 3.967 | 4.246 |
| 8 | UK | Social | CV | - | - | 24.103 | 4.319 | 4.078 |
| 8 | UK | Social | SCI | 0.25 | - | 26.095 | 5.315 | 4.512 |
| 8 | UK | Social | QT | 0.2 | 0.02 | 26.528 | 4.941 | 4.229 |
| 12 | UK | Social | CV | - | - | 21.439 | 6.361 | 4.852 |
| 12 | UK | Social | SCI | 0.4 | - | 21.878 | 8.634 | 6.737 |
| 12 | UK | Social | QT | 0.1 | 0.04 | 22.881 | 7.695 | 6.544 |
| 18 | UK | Social | CV | - | - | 25.145 | 8.261 | 5.671 |
| 18 | UK | Social | SCI | 0.2 | - | 26.392 | 8.209 | 6.627 |
| 18 | UK | Social | QT | 0.2 | 0.005 | 27.195 | 7.624 | 6.136 |
| 24 | UK | Social | CV | - | - | 25.747 | 6.292 | 4.719 |
| 24 | UK | Social | SCI | 0.1 | - | 26.339 | 7.128 | 5.891 |
| 24 | UK | Social | QT | 0.1 | 0.015 | 26.577 | 6.976 | 6.145 |

*Figure S 5*

### Supplementary Materials 5: Effect on data quality and retention of QT-NIRS thresholds

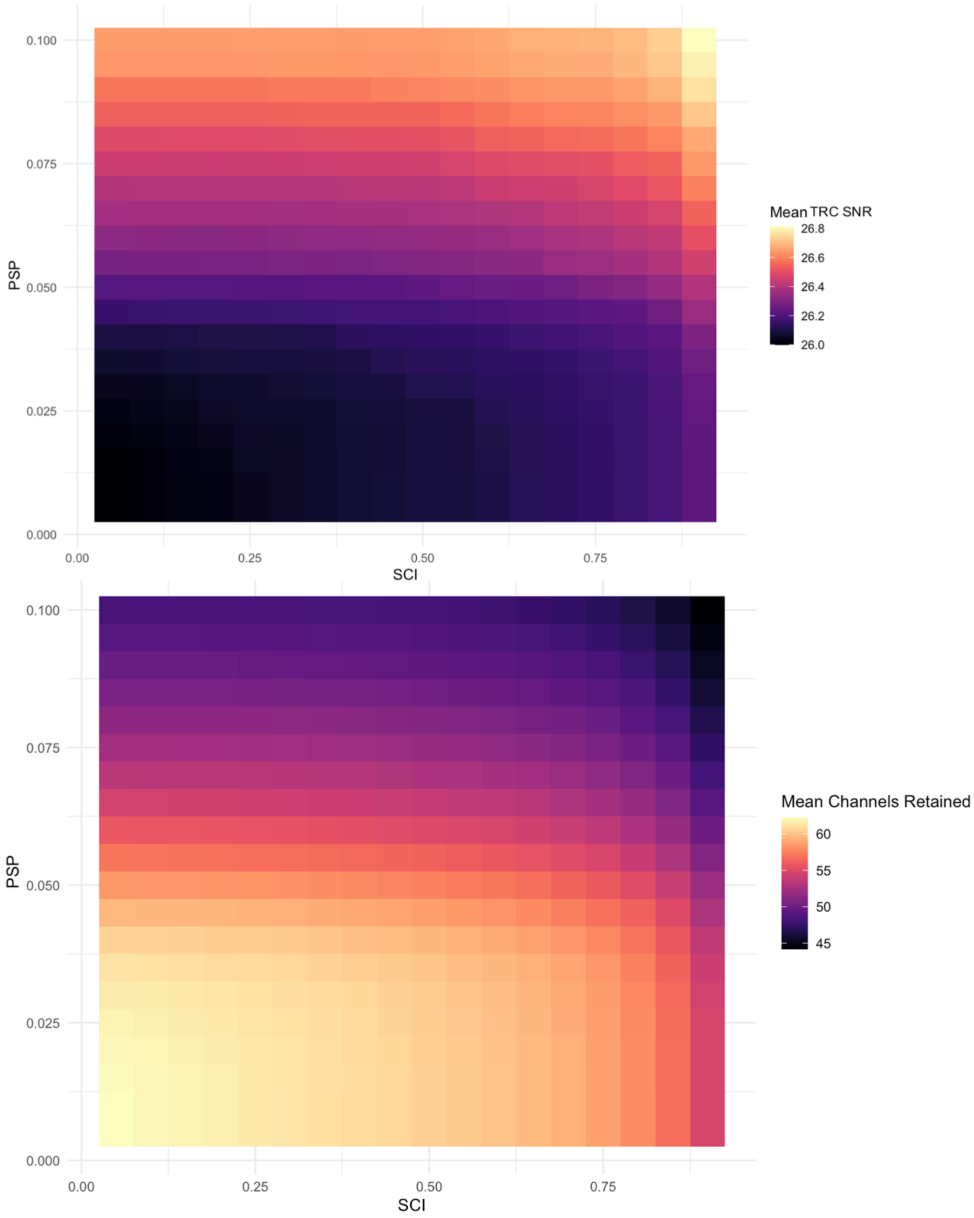


*Figure S 6: Heatmaps showing fNIRS signal quality and retention for each SCI- and PSP Threshold value combination.*

*Top: TRC SNR for each combination of parameter values, showing higher/lower values corresponding for combinations incorporating high/low values for both parameters, respectively. Bottom: Channels Retained for each combination of parameter values, showing higher/lower numbers of channels retained for combinations incorporating low/high values for both parameters, respectively.*

### Supplementary Materials 5: Relationship between data quality and age

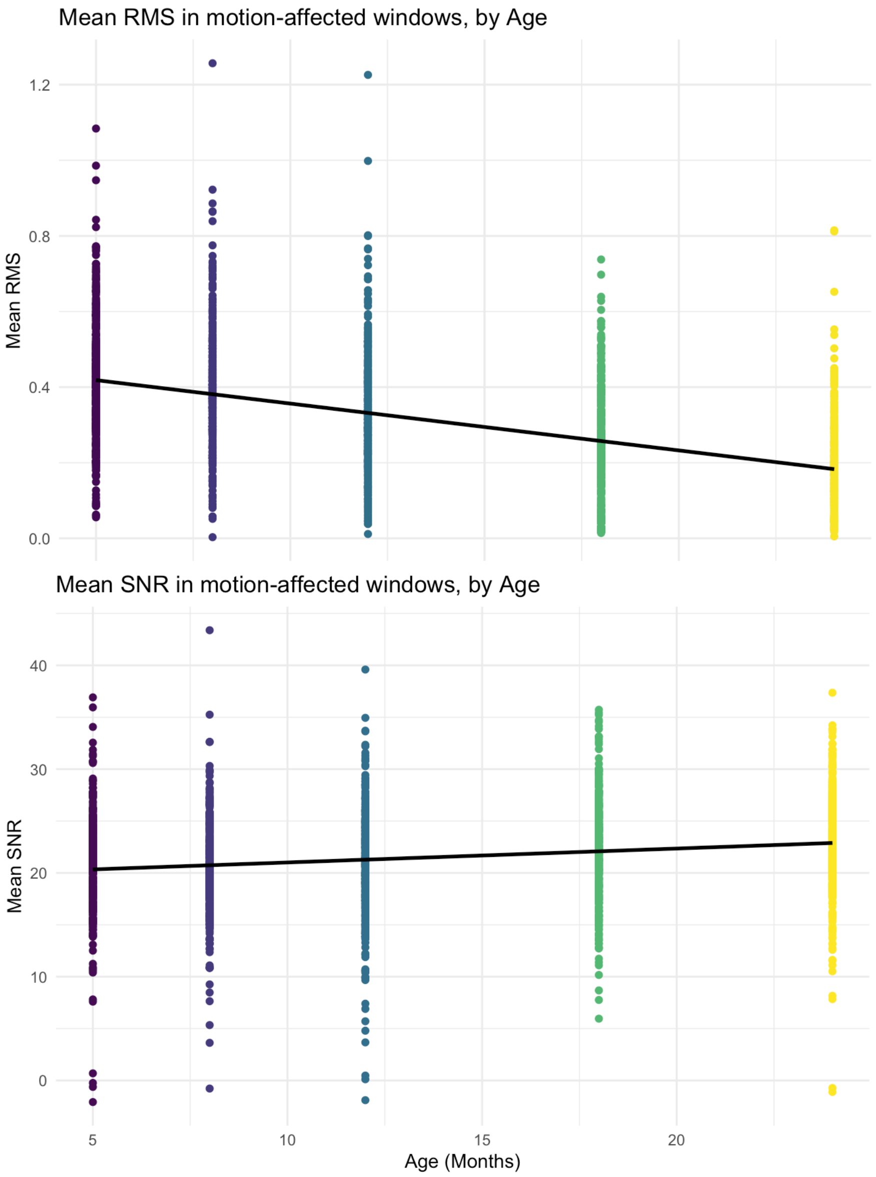


***Figure S 7: Motion severity in detected artefact windows, by age***

*Top: channel-wise means of root mean squared values within windows containing detected artefacts. Bottom: channel-wise means of signal-to-noise ratio values within windows containing detected artefacts.*
